# Discovery of *Mycodnaviridae*, a clade of giant viruses that persistently infect zoosporic fungi

**DOI:** 10.1101/2024.01.04.574182

**Authors:** Jillian M. Myers, Frederik Schulz, Saleh Rahimlou, Vikas Yadav, Sheng Sun, Kevin R. Amses, D. Rabern Simmons, Michelle Orozco-Quime, Joseph Heitman, Jason E. Stajich, Timothy Y. James

## Abstract

Giant viruses of the phylum *Nucleocytoviricota* have emerged as particularly notable due to their increasingly recognized impacts on eukaryotic genome evolution. Their origins are hypothesized to predate or coincide with the diversification of eukaryotes, and they have been detected in hosts that span the eukaryotic tree of life. But surprisingly, such viruses have not been definitively found in Kingdom Fungi, though earlier genomic and metagenomic work suggests putative associations. Here we report both “viral fossils” and active infection by giant viruses in fungi, particularly in the zoosporic phyla Blastocladiomycota and Chytridiomycota. The recovered viral assemblies span up to 350 kb, encode over 300 genes, and form a monophyletic family-level clade within the *Nucleocytoviricota* related to orders *Imitervirales* and *Algavirales*, which we name *Mycodnaviridae*. We observed variation in infection status among the isolates including apparent active infection and transcriptionally suppressed states, suggesting that viral activation may be constrained to certain life stages of the host. Our experimental findings add to the limited natural virus-host systems available in culture for the study of giant viruses and expand the known host range of *Nucleocytoviricota* into a new kingdom that contains many model species. *Mycodnaviridae* have a global distribution, which invites inquiry into the implications of these infections for host traits, host genome evolution, and the metabolic impacts on ecosystems.

## Main

In the entire Fungal Kingdom, only a handful of DNA viruses are known–all small circular single-stranded (CRESS) viruses (Li 2020; Yu 2010). This apparent rarity of DNA viruses in fungi likely results from taxonomic biases in searches, rather than a true biological phenomenon. The study of mycoviruses has historically focused on their application for biological control of fungal pathogens, particularly in agricultural contexts, and as a result most research has involved viruses infecting fungi in the Ascomycota—one of two phyla comprising the subkingdom Dikarya, which contains over half of described fungal taxa. The current fungal taxonomic division includes eight phyla of zoosporic fungi that reproduce with motile spores and represent a particularly understudied paraphyletic assemblage that have retained ancestral characters shared with animals (Prostak 2021, Hyde 2024). We previously demonstrated that by expanding the taxonomic scope of considered fungal hosts to include these lineages outside of the Dikarya the known RNA mycoviral diversity was greatly increased (Myers 2020). However, the presence of DNA viruses in these fungi has been almost entirely unexplored (but see Clemons 2023).

Indeed, the finding of giant viruses (GVs), which we report here, began as a serendipitous discovery resulting from a large genomic sequencing effort across these fungal phyla (Amses 2022).

Nucleocytoviricota is a phylum of highly diverse “giant viruses” including those with the largest and most complex genomes yet known, which is a result of gene acquisition from eukaryotic hosts and bacteria as well as substantial gene duplication in some lineages (Koonin and Yutin 2018). Since their discovery GVs have challenged the established expectations in virology as some encode genes involved in transcription and translation (Raoult 2004, Schulz 2017, Fels 2026) as well as various metabolic processes (Monier 2017, Schvarcz 2018, Blanc-Mathieu 2021). Studies have shown that up to 10^^^5 GV genomes can exist within a single microliter of ocean water (Hingamp 2013) and the taxon richness of just one order, *Imitervirales*, surpasses that of both bacteria and archaea in oceans (Mihara 2018).

Despite their incredible abundance, relatively few have been linked to a host, and only a scant number of natural GV-host systems have been studied in the laboratory. Much of what is known about giant virus biology has been learned from protist-GV interactions. Lytic dynamics are common wherein following endocytosis of the virus into the host cell the viral DNA is transported to the nucleus for replication, then to cytoplasmic viral factories for assembly, and virions exit the cell by budding or cell lysis. Interestingly, some GVs in the Algavirales can also enter a latent phase wherein the viral genome is integrated into a host chromosome. Reactivation of the proviruses can be triggered by temperature, but the molecular mechanisms involved are unknown (reviewed in Duchêne 2026). GVs must certainly have meaningful ecosystem-level biogeochemical consequences which have yet to be fully understood or quantified. The development of additional laboratory model systems of GVs and their natural hosts could lead to advances in our understanding of virus-host interactions, including host immune response, viral “reprogramming” of host cells, metabolic impacts, and their ecosystem-level consequences.

Confirmed associations with GVs include hosts from a breadth of taxa—from single-celled algae to invertebrates to animals (reviewed in Koonin 2015)—and metagenomic- and metatranscriptomic-powered inferences suggest an even broader scope of eukaryotic hosts (Moniruzzaman 2017, Schulz 2020). Previous studies have suggested associations between *Nucleocytoviricota* and fungi through the identification of putative fungal homologs in metagenomic-assembled giant viruses (Schulz et al. 2018, Schulz et al. 2020, Bhattacharjee et al. 2023) and giant virus EVEs in fungal genomes (Gallot-Lavallée and Blanc 2017, Gong et al. 2020, Zhao et al. 2023, Nobre 2024). Of these mentioned, only the EVE studies directly connect fungi with viruses, while the metagenomics studies predict fungal affiliation of the viruses based on gene homology and, in the case of Bhattacharjee et al. 2023, the dominance of fungi in the environment. The latter does not preclude the possibility of non-fungal hosts. Active GV infections in fungi have not previously been documented nor have novel clades that include active giant viruses in fungi been characterized. The cumulative evidence suggests a rich history between GVs and fungi, with much more yet to be understood.

## Results

### Zoosporic fungi contain giant virus genes and genomes with global distribution

We identified homologs of the giant virus major capsid protein (MCP; NCVOG0022) in the genomes of 17/141 fungi with HMM searches (Fig S1, Table S1). The distribution of MCP homologs is restricted to non-Dikaryotic lineages, and, particularly, the zoosporic fungi; no MCPs were found by hmmsearch in non-zoosporic lineages (n= 40). We further searched all fungal genomes in Mycocosm (Grigoriev 2014; n = 2,543) for MCP homologs by blastp and protein model searches, which resulted in one hit to a non-zoosporic fungal species: *Dimargaris cristalligena* (phylum Zoopagomycota), a mycoparasite. Among the zoosporic lineages, MCPs were found in Chytridiomycota (n=12), Monoblephidomycota (n=2), and Blastocladiomycota (n=3). Blastocladiomycota are a particularly poorly sampled group, and thus our genomic searches were limited (n=7). However, MCPs appear to be common in the most highly sampled genus within this phylum, *Allomyces*. Based on available sequences, we designed primers targeting the MCP gene of *Allomyces* and found 30/58 unique isolates were positive for capsid by PCR.

The contigs containing MCP genes vary widely among the fungal isolates in gene content, GC composition, intergenic distance, and other metrics, suggesting numerous independent endogenization events (Fig 1 A). In three isolates (*Podochytrium* sp. JEL797, *Gaertneriomyces semiglobifer* Barr43, and *Fimicolochytrium jonesii* JEL569), MCP was the only gene of viral provenance on contigs which otherwise contain primarily fungal genes and have GC content and intron density similar to other host contigs (Fig S2). These MCPs may be the result of horizontal gene transfer (HGT) from GVs, or remnants from past latent GV infections. In other isolates, contigs with MCPs appear as mosaics of viral, prokaryotic, and fungal genes, which include multiple additional giant virus orthologous genes (GVOGs), and have markedly greater gene density compared to host contigs, consistent with reference GV genomes (Fig 1 A & B; Fig S2). These giant virus-like contigs range up to nearly 350 kb and are suggestive of either very recent endogenization or, as we later evidence, proviral (latent) state. Interestingly, in five isolates the MCP contig contains a gene encoding an integrase with sequence similarity to those of polinton-like viruses (*Eupolintoviridae*). Polintons are dsDNA transposons of eukaryotes with ancient and enigmatic associations with GVs (Koonin 2024). Our finding of polinton-like integrases coinciding with nucleocytoviricot-like polB and other GVOGs may help elucidate the relationships of these groups.

**Fig 1.**
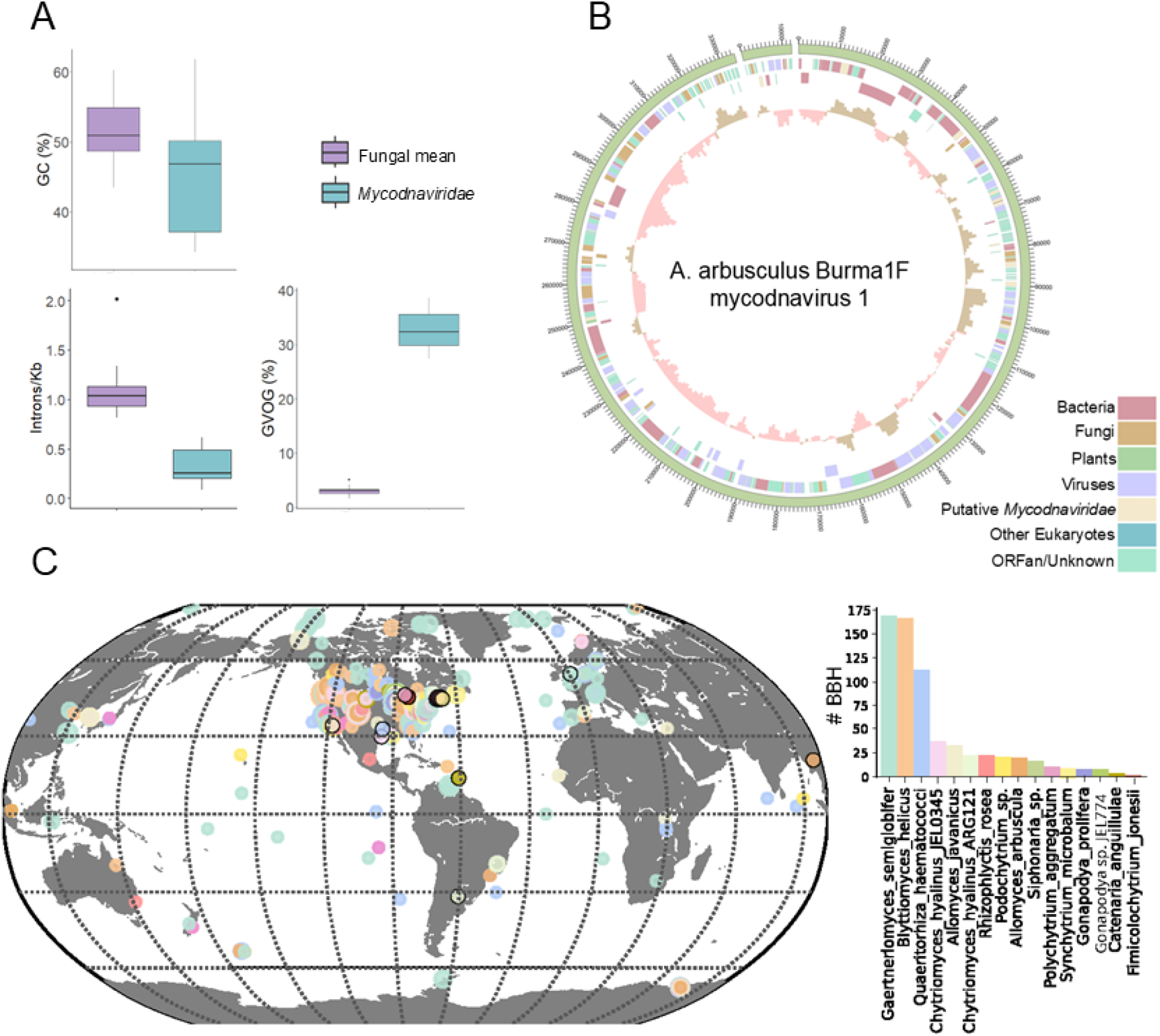
Genomic features of *Mycodnaviridae*. (A) Boxplots of genome statistics for the fungal genomes found to contain MCP genes (n=17): mean values across all contigs (“Fungal mean”, purple), and MCP contigs that met the criteria set for orthology analysis (ViralRecall score >2, viral region length > 70%, intron density >25% different from host; “*Mycodnaviridae”*, teal). (B) The genome of Allomyces arbusculus Burma1F mycodnavirus 1 is a mosaic of bacteria; viral, and eukaryotic genes. Contigs are represented by the outermost ring in the circularized genome plot. From outside to inside: ORFs on the forward strand, ORFs on the reverse strand (both color-coded by provenance), GC skew. (C) Global distribution of the *Mycodnaviridae-*like sequences based on the major capsid protein. Colored circles correspond to metagenomic MCP sequences with best blast hit (BBH) to *Mycodnaviridae* identified in this study and circles with a black outline indicate geographic location of *Mycodnaviridae* and hosts described in this study.

Based on MCP data previously extracted from global metagenomics datasets, we assessed the global distribution of these novel fungal MCPs. Environmental MCP homologs with bidirectional best blast hits to fungal MCPs have a global distribution (Fig 1 C). The *Allomyces* isolates tested by PCR for MCP have worldwide provenance, thus also suggestive of a long history of viral infection.

We first closely analyzed the GV genome assemblies from two strains of *Allomyces*. A. arbusculus Burma1F mycodnavirus 1 includes two contigs of total size 345,111 bp, encoding 314 genes with likely provenance, based on homology, assigned as follows: 10% fungi, 14% bacteria, 20% virus, 4% putatively *Mycodnaviridae-*specific, and 52% unknown/ORFan (Fig 1 B). The isolate *A. javanicus* California12 has multiple large contigs (ranging between 80-288kb) with giant virus signatures (Fig S3). Interestingly, when binned by tetramer frequency, the viral contigs in *A. javanicus* California12 form two bins suggesting the presence of two distinct viruses. The two bins are roughly similar in size (212,591 bp and 287,844 bp) and show some duplication of hallmark GVOGs: each bin contains copies of MCP, D5-like helicase-primase, DNA PolB, mRNA capping enzyme large subunit, and superfamily II helicase. The larger bin contains an additional five GVOGs (YqaJ recombinase, RNA ligase, disulfide (thiol) oxidoreductase, transcription initiation factor IIB, and A32-like packaging ATPase). *A. javanicus* strains are believed to be hybrids (Emerson 1941) and the two viruses could have originated from the two parents of this diploid sporophyte. Alternatively, this finding could be a result of co-infection by two distinct virus species.

### *Mycodnaviridae* is a monophyletic clade of fungal giant viruses

An updated species tree of representative *Nucleocytoviricota*, based on a concatenated alignment of the 7 protein GVOG7 (Aylward 2021) showed that five of the 12 fungal-associated EVEs contained sufficient marker recovery (>=3/7 GVOGs) for inclusion in the final tree. These genomes formed a monophyletic clade with the three metagenome-assembled genomes (GVMAGs) previously assigned to IM_02, indicating that the former IM_02 lineage is expanded and here treated as MY_02 or *Mycodnaviridae* (Fig 2 A; Vasquez 2025). In the revised phylogeny, *Mycodnaviridae* is recovered within a broader order-level clade basal to *Imitervirales*, consistent with the taxonomic framework proposed by Vasquez et al. (2025). This supports the main conclusion that fungal giant viruses cluster with the former IM_02 lineage. Hosts of this lineage are unknown, but the clustering of *Mycodnaviridae* in this clade may support a fungal association for the other GVMAGs in MY_02.

**Fig 2.**
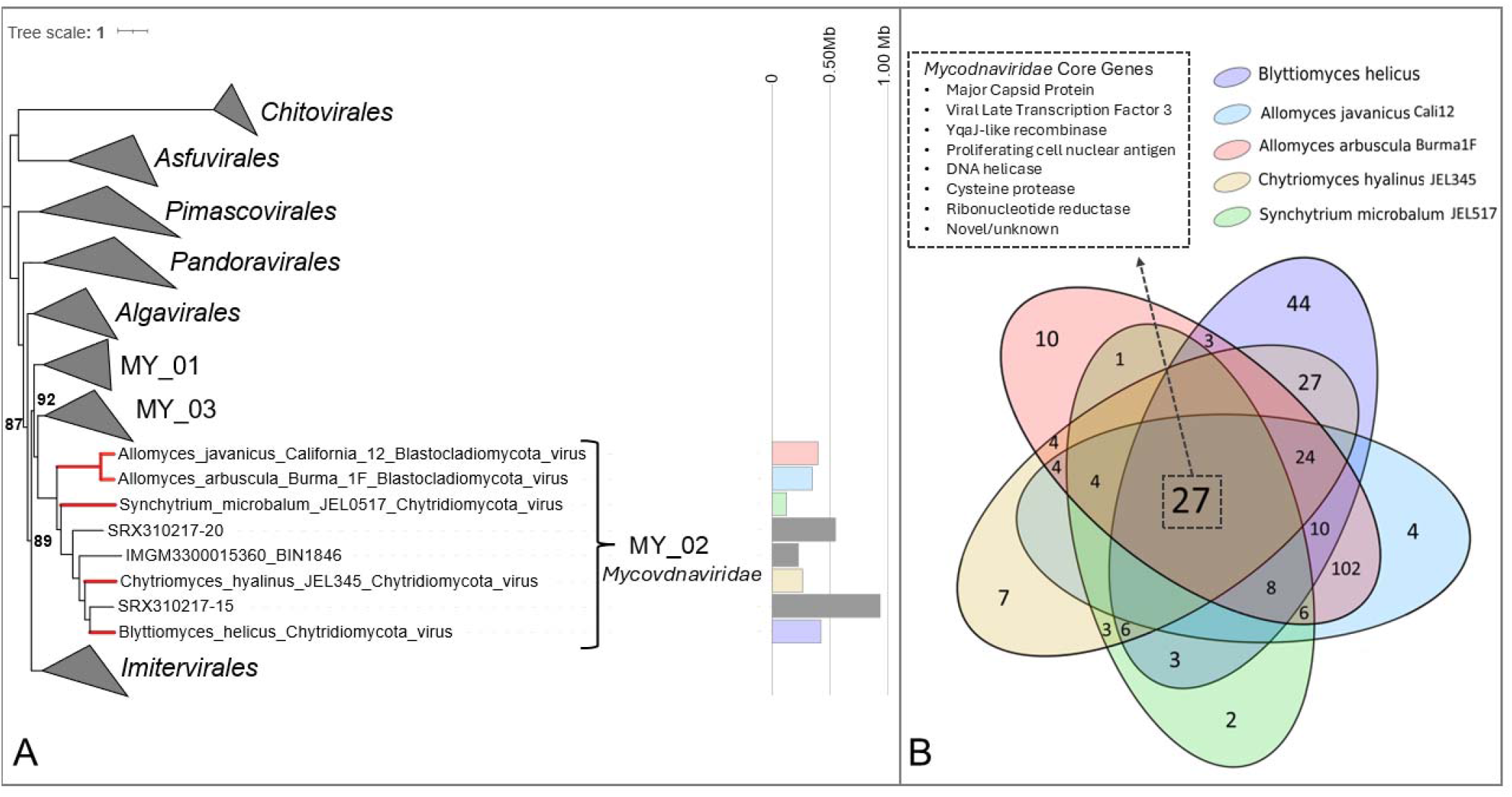
Phylogenetic position of the *Mycodnaviridae* and core genes. (A) Position of the *Mycodnaviridae* clade in the species tree of the *Nucleocytoviricota*, wherein the un-collapsed branches represent *Mycodnaviridae*, and MY_01-03 are as in Vasquez 2025; within this clade, branches highlighted in red indicate those genomes sequenced from fungi, and branches in black indicate metagenomics-assembled genomes previously known as IM_02. The tree is rooted at Pokkesviricetes. Bars to the right indicate recovered assembly sizes. (B) Venn diagram indicating the number of orthologous gene clusters shared between the 5 most complete *Mycodnaviridae* genomes.

We investigated whether a core suite of genes exists that functionally define the *Mycodnaviridae* by conducting orthology analysis on the most complete GV assemblies–five have at least 9 of the 20 core GVOGs present in the majority of the *Nucleocytoviricota* (Yutin 2009). This resulted in 27 gene clusters shared by all five (Fig 2 B) though this is likely an underestimate due to variation in assembly completeness, which is suggested by the variation in GV assembly sizes recovered from different fungal hosts. In addition to *Nucleocytoviricota* core GVOGs such as MCPs, VLTF3, and YqaJ-like recombinase, the *Mycodnaviridae* core set is composed of gene orthogroups involved in DNA biosynthesis such as DNA helicase, ribonucleotide reductase, and proliferating cell nuclear antigen, which are also GVOGs found in other large and giant viruses. Other shared gene clusters include a cysteine protease, protein with N-acetylglucosaminyltransferase activity (possibly involved in cell wall modification), six genes of unknown function, and five novel genes apparently unique to these viruses.

### *Allomyces* viruses are integrated in the nuclear genome and show variation in methylation modification patterns

Critical questions about the giant virus elements are where the apparently complete elements are in the genome and whether and when they are capable of replicative infection. We conducted long-read whole-genome sequencing by Oxford Nanopore to determine whether the viral contigs previously recovered are contiguous in the host genome and to detect methylation modification. We also conducted RNA-Seq to assess expression of viral genes and quantitative PCR of MCP genes as a proxy for viral titer.

In *Allomyces arbusculus* Burma1F, long-read sequencing revealed the viral contigs side-by-side and integrated into a chromosome that harbors telomere repeats on both ends (contig 03: 1397320-1743578; Fig 3 A & B). Nanopore-based genome assembly also revealed 13 clusters of LTR-transposons, only one per contig, that potentially represent centromeres similar to those observed in the dikaryotic fungus *Cryptococcus neoformans* (Yadav 2018, Janbon 2014). With interest in whether the viral genes are expressed and, if so, whether expression occurs throughout host tissue or is constrained to some structures or life cycle stages, we sequenced RNA of *A. arbusculus* Burma1F at three different stages. We found almost no evidence of expression from the viral genes, with the exception of genes in the flanking region (Fig 3 B). CpG methylation was not detected in these viral regions per the Nanopore-generated data but, curiously, 6mA methylation was reduced relative to the rest of the chromosome (Fig 3 B). Thus, while the expression of A. arbusculus Burma1F mycodnavirus 1 was apparently silenced at the time of sequencing, the mechanism is as yet unknown.

**Figure 3.**
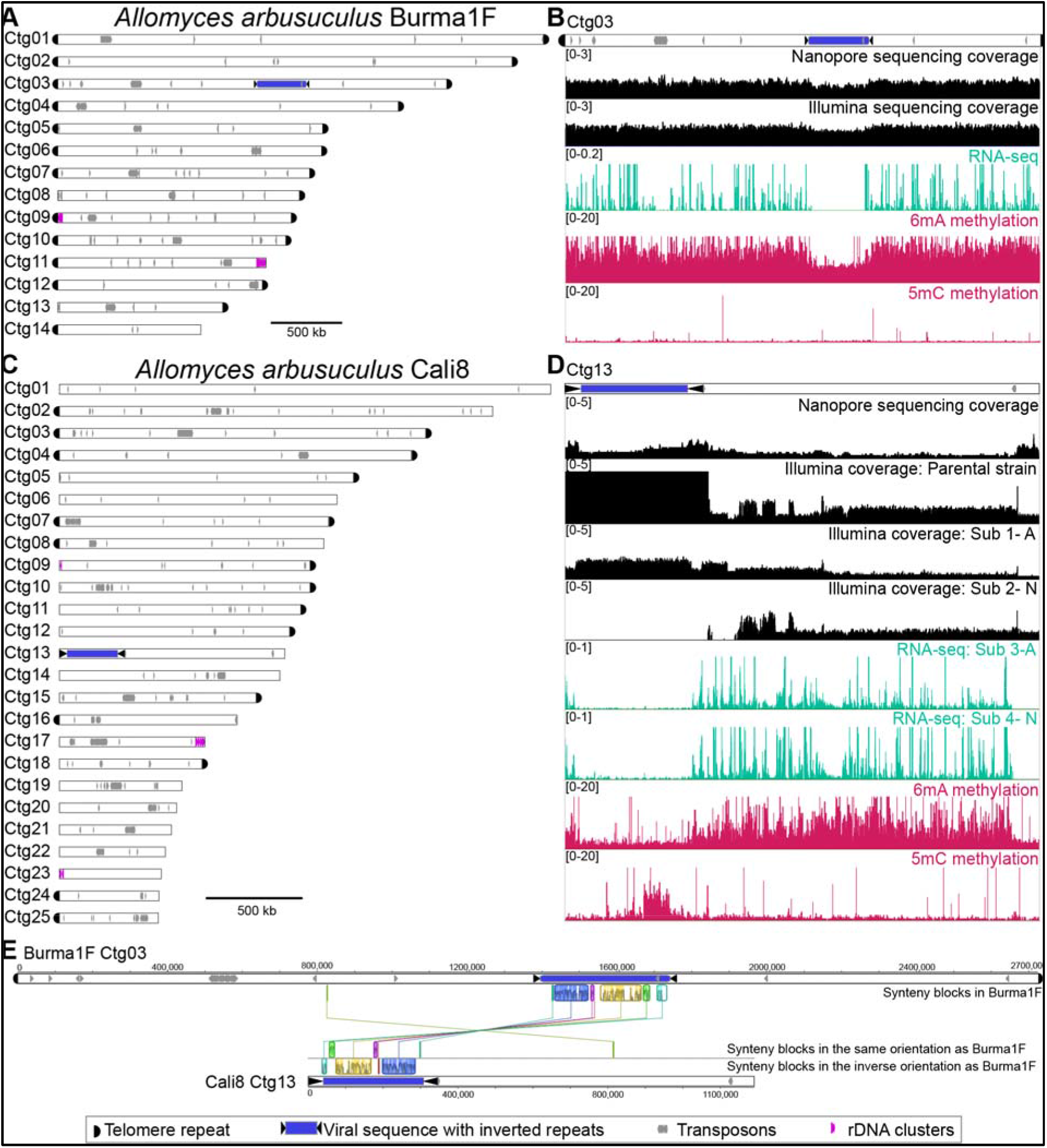
*Allomyces* viruses are integrated into the nuclear genome, with complex patterns of expression/suppression and methylation modification. (A) Nanopore-based assembly of *A. arbusculus* Burma1F shows the virus integrated into the host nuclear genome (Ctg 03, blue) (B) Mapping of Nanopore reads, Illumina sequencing reads, RNAseq reads, 6mA and CpG methylation extracted from Nanopore sequencing reads across the virus-harboring contig of *A. arbusculus* Burma1F. Note that the copy number of the viral region for JM-6 exceeds the y-axis displayed, see Fig S4 (C) Nanopore-based assembly of *A. arbusculus* Cali8 shows the virus integrated into the host nuclear genome (Ctg 13, blue). (D) Mapping of Cali8 Nanopore reads, Illumina sequencing reads of original strain (“Parental strain”) and strains with abnormal (“Sub 1-A”) and normal phenotype (“Sub 2-N”), RNAseq reads of a Cali8 strain with abnormal (“Sub 3-A”) and normal phenotype (“Sub 4-N”), and 6mA and CpG methylation extracted from Nanopore sequencing reads across the virus-harboring contig of *A. arbusculus* Cali8. (E) Viral insertion regions of B1F and Cali8 are largely not syntenic, while the viruses themselves are highly syntenic with regions in inverted orientation. In (B) and (D), the bracketed numbers denote the y-axis scale. Nanopore and Illumina sequencing data are shown as copy number (read coverage normalized by average genome coverage), RNA-seq data is shown as RPKM (normalized per-base coverage as bins per million reads), and methylation data as fraction of reads methylated.

We performed both short- and long-read sequencing of an additional *Allomyces* strain, *A. arbusculus* Cali8, and observed a different pattern. Though the Illumina assembly was more fragmented for Cali8 than Burma1F, viral contigs were detected that harbored MCPs, seven additional core GVOGs, and a viral-like integrase. Interestingly, these contigs averaged nearly 50-fold higher coverage than the rest of the genome (Fig 3 D, “Parental Strain”; Fig S4), suggesting that these DNAs were in high titer at the time of sequencing. When mapped onto the nanopore-based assembly, these high-coverage contigs were side-by-side (Fig 3 C), and flanked by additional high coverage regions on either side for a total GV assembly size of approximately 350kb. Like Burma1F, 6mA methylation was detected across the viral region at a lower rate than the rest of the genome (Fig 3 D); but in contrast to patterns in Burma1F, CpG methylation was detected in parts of the viral region (Fig 3 D).

We compared the host genomic regions of viral sequence integration in Cali8 versus B1F. We found only one high-confidence syntenic region >1kb (Fig 3 E). Both endogenous viruses harbor inverted terminal repeats. Within a species, these repeats were > 99% identical on either side, however the sequences are not conserved between species. The repeats on the termini of A. arbusculus Burma1F mycodnavirus 1 are approximately 9kb, while the repeats on the termini of A. arbusculus Cali8 mycodnavirus 1 are an astonishing 34kb. The viral regions themselves overall are syntenic, although seem to have undergone significant structural rearrangements with large syntenic blocks in the inverse orientation between the two isolates (Fig 3 E).

### Evidence for viral replication and disease phenotype

Our initial clue that viral replication was occurring in Cali8 was based on high coverage of viral contigs in our short-read sequencing data. We reisolated this strain from our stock of resting sporangia and attempted to trigger viral replication, assessed the morphology for evidence of a diseased phenotype, re-sequenced the genomes, performed RNA-Seq, and conducted quantitative-PCR assays to assess viral gene abundance.

Fungi in the phylum Blastocladiomycota, such as *Allomyces*, have life cycles that include the alternation of generations as haploid gametophytes (which produce haploid gametes) and as diploid sporophytes (which produce diploid spores) (Fig S5). During the sexual phase of the typical healthy culture of *Allomyces* a haploid meiospore germinates to produce a gametophyte with pairs of “male” and “female” reproductive structures (gametangia), each producing gametes which fuse to ultimately form a diploid sporophyte. The mature sporophyte can continue growth in a diploid asexual cycle and also produce resting sporangia that undergo meiosis to produce haploid meiospores which develop into gametophytes after a brief motile period. We found that gametophytes of Cali8, unlike those of Burma1F and Cali12, showed a high rate of mortality and morphological and growth rate variation (Fig 4 A). Some gametophytes had typical growth and development with terminal female gametangia above orange pigmented male gametangia (Fig 4 B). Other strains showed varying degrees of abnormal development, but often including atypical “bloated males”, with pigment and discharge papillae like male gametangia, but not adjacent to female gametangia as in normal gametophyte development (Fig 4 B). These bloated males failed to release gametes. Unusual twisted hyphae and blebbing at hyphal tips were also observed, and this abnormal phenotype strain showed a reduced growth rate (Fig 4 C & D).

**Figure 4.**
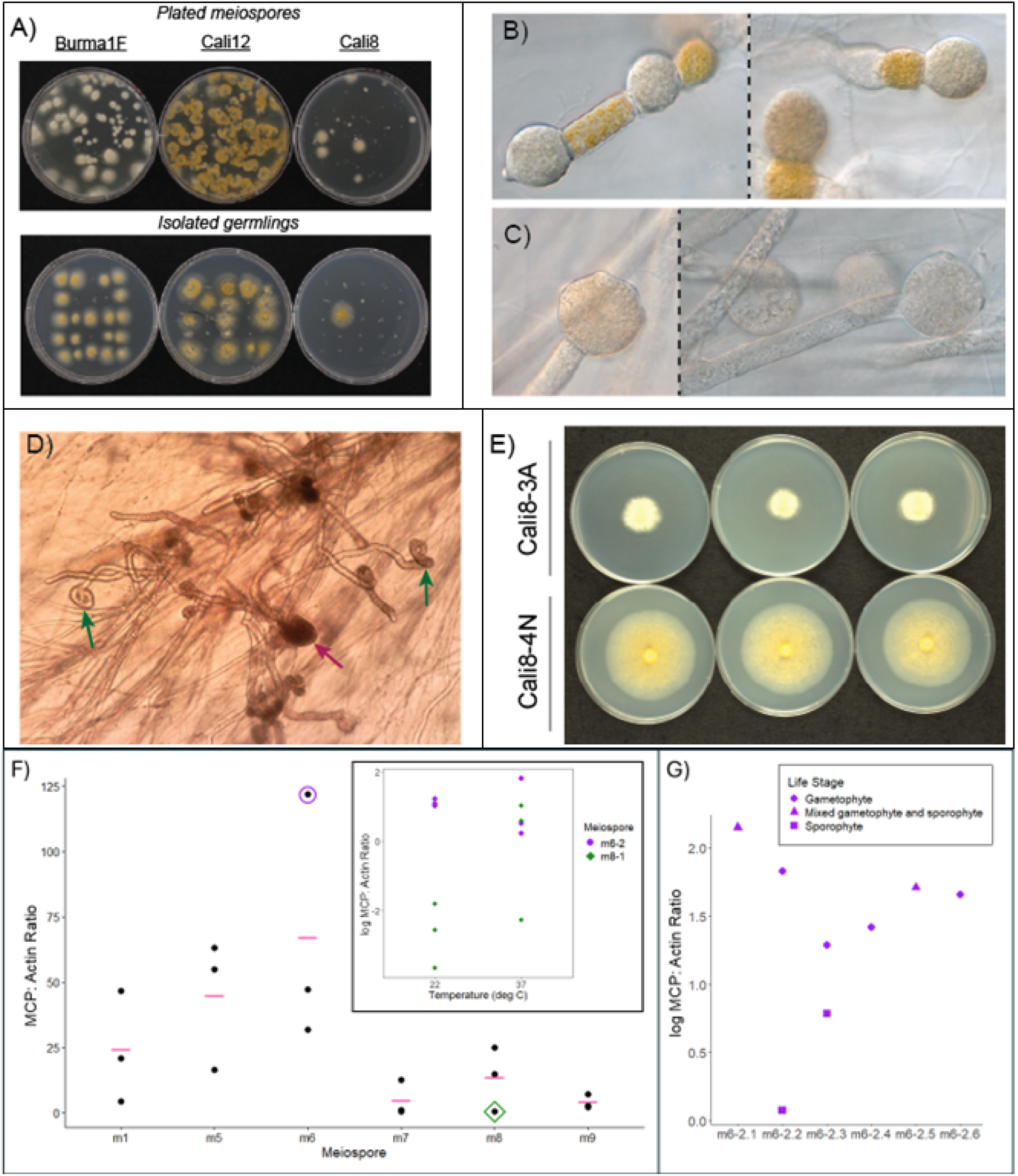
(A) Meiospores of three *Allomyces* isolates show differences in (B) Cali8-N subcultures display normal gametophyte phenotype (C) Cali8-A subcultures display malformed gametophytes with swollen and unproductive male gametangia (D) Unusual twisted hyphae (green arrows) and blebbing (pink arrow) in Cali8-A (E) Cali8-A has a reduced growth rate relative to Cali8-N. Bottom panel shows qPCR results as the ratio of MCP to actin DNA copies, a proxy for viral abundance. (F) Isolates grown from individual meiospores from the same parent varied significantly (p <.001) in MCP:actin ratio (individual data points are shown as black dots, means are shown as pink bars). An isolate with very high viral abundance (m6-2, purple circle) and very low viral abundance (m8-1, green diamond) were subcultured at 22C and 37C (inset). (G) M6-2 was further subcultured and MCP:actin ratio assessed at various life stages.

We then investigated the association of these phenotypes with presence of the virus using genome sequencing and qPCR. We sequenced the genomes of two Cali8 gametophyte cultures, one of which had the abnormal phenotype (Sub-culture 1-A) and the other of which had presumably selfed and had sporophyte morphology (Sub-culture 2-N). The abnormal Sub1-A had higher coverage of the viral region compared to the rest of the genome, though not as high of coverage as the originally sequenced strain (Fig 3 D). The strain with normal sporophyte phenotype (Sub2-N) had almost no sequencing coverage of the viral region (Fig 3 D), suggesting that the virus had been lost. We sequenced RNA from two additional subcultures which had distinct growth profiles (Fig 4 D), displaying either diseased (Sub-culture 3-A) or normal (Sub-culture 4-N) phenotype. Surprisingly, for both subcultures the RNA-seq data show similar, very low expression of the viral region (Fig 3 D). While these results are complicated, together they suggest that in the Cali8 strain the virus can be expressed though it does not consistently result in a diseased phenotype. These results also suggest that in some cases the *Allomyces* host is able to rid itself of the virus.

One mechanism that could result in virus genome disruption or elimination is meiosis. qPCR demonstrated that the gametophytes derived from different meiospores varied significantly in viral titer (ANOVA F-statistic = 3.36559, p= 0.03944) (Fig 4 E).Variation is highly dynamic, and multiple subcultures of the same gametophyte produced differing amounts of virus. While qPCR of Cali8 cDNA showed that the virus is typically not expressed, MCP cDNA did amplify in one Cali8 gametophyte sub-culture, indicating expression of this viral gene. qPCR of cultures grown at high temperatures showed a greater variance in viral titer, suggesting that heat can trigger viral replication (Fig 4 F). The unreliability of heat as a viral trigger indicates that other factors, such as other environmental conditions, host density, or life cycle stages, are also involved. Using both filtered cell homogenate and supernatant of a Cali8 subculture with abnormal morphology (Sub-cultures m1 and m1-3), we attempted to infect the virus-free strain Mex58 by applying these crude extracts to mating gametes but were unsuccessful.

These results strongly indicate an active infection by a giant virus in this strain of *Allomyces*. Other sequenced *Allomyces* strains had contigs similarly rich with GV hallmark genes but lacking differences in coverage from the genome-wide median estimated by read depth. It is plausible that in these strains the conditions for triggering viral release were not met under laboratory culture conditions, and thus the virus remained in a proviral state. Given the high occurrence of MCPs we suspect that additional *Allomyces* strains contain provirus EVEs that are capable of activating and forming an infectious propagule, though the triggering mechanism for viral production is unknown.

## Discussion

We report the discovery of *Mycodnaviridae*, a new family of *Nucleocytoviricota* with close phylogenetic affinities to *Algavirales* and *Imitervirales*. The three phyla of fungi in which we have found evidence of giant viruses—Blastocladiomycota, Chytridiomycota, and Zoopagomycota— have life cycle stages similar to some algae and amoebae, which may facilitate infection by viruses with persistent infection strategies, such as the phaeoviruses in *Algavirales*. In both algae and zoosporic fungi there exist life-stages (spores and gametes) consisting of flagellated single-cells lacking a protective cell wall. The phaeoviruses have evolved stable coexistence with their hosts through a persistent viral strategy whereby these spores or gametes are infected by virus particle(s) that then integrate into the host genome (Müller and Parodi 1993). Development proceeds with one copy of the virus integrated into the DNA of each cell (Meints 2008). Thalli often appear symptom-free until some unknown mechanism triggers virus emergence from the genome. Reproductive organs may develop irregularly as they become factories for viral replication (Henry and Meints 1992; Stevens 2014). Very recently, another integrated viral genome was identified within a green alga and was shown to actively produce viral particles (Erazo-Garcia 2025). We hypothesize that fungal giant viruses in zoosporic lineages may share similar replication cycle characteristics wherein life cycle stages lacking a cell wall may be infected and then endogenized into the genome during the vegetative stages where the host proliferates with a cell wall (Fig S5).

Genome coverage and qPCR data, transcriptional patterns, and the presence of an integrase gene in *Mycodnaviridae* are consistent with this hypothesized replication cycle. In most genomes viral contigs have similar coverage to the host average, as would be expected for a provirus integrated into a single-celled spore/gamete that propagates the organism. Genome coverage data in one isolate of *Allomyces* (Cali8), however, suggests active replication at the time of initial sequencing. Gametophytes developed from individual meiospores of the same culture show differing amounts of MCP DNAs per qPCR, and these differences are inducible by altering the growth temperature. Though speculative, this could be indicative that, like the phaeoviruses, these reproductive structures may become GV replication factories under some conditions. Gametophyte qPCR indicated near loss of the virus in some individuals; like the phaeoviruses, it seems possible that A. arbusculus Cali8 mycodnavirus 1 may be lost through meiosis. Sporophytes of this isolate produce fewer MCP per qPCR compared to gametophytes or mixed cultures, consistent with endogenization at this life stage. While RNA-seq of Cali8 showed only slight expression of the viral region, this is an interesting departure from the complete lack of expression observed in Burma1F. Though we sequenced RNA at multiple life stages in Burma1F, we unfortunately did not capture RNA from the gametophyte stage, which may explain why viral activation was not observed in this isolate. The observed transcriptional patterns in Burma1F suggest that the provirus may be silenced in the host genome, also consistent with transcriptional patterns of phaeoviruses integrated in their algal hosts (Cock 2010). Further investigation in this strain may reveal conditions under which the virus is activated. Future studies should also address whether, like the phaeoviruses, the virus genome can circularize. These results highlight that viral release is complicated and may be linked with environmental and host lifecycle triggers as well as the complex life history of this fungus. These results also indicate that the host possesses mechanisms for avoiding viral release and/or restricting the spread of viral particles.

Like all cellular organisms, the evolution of the fungi has been impacted by interactions with viruses. Giant viruses are hypothesized to have originated with the diversification of eukaryotes, and metagenomic sequence data demonstrates their abundance in the environment. The lack of evidence of active infection by giant viruses in fungi before now is therefore surprising, though previous reports have shown their association (Zhao 2023; Bhattacharjee 2023; Gong 2020; Schulz 2020; 2018). Most interestingly, the EVEs reported by others are of distinct and distant relation to the clade of giant fungal viruses we report here. Phylogenetic reconstruction of the DNA polymerase B (PolB) using data from four such studies as well as *Mycodnaviridae* shows at least five unique clades of giant virus PolBs associated with Fungi (Supplemental Figure S6). Taken together, these data suggest that, like animals, fungi have historically and, perhaps, currently been hosts to multiple distinct lineages of giant viruses.

It is intriguing to consider the implications of a possible ancient association between fungi and giant viruses in relation to fungal evolution because viral gene acquisition can be an impetus for evolutionary innovation (Frank and Feschotte 2017). In the fungi, a transcription factor involved in cell cycle regulation that is, surprisingly, unique to fungi is hypothesized to have been acquired by the fungal ancestor through lateral gene transfer from a virus (Medina 2016). Phylogenetic evidence points to the source as either a giant virus or phage-infected bacterium. We searched MCP-containing contigs but found no homologs containing the diagnostic viral-like KilA-N domain of this cell-cycle protein, though this does not preclude giant viruses as the original source. Other evolutionary innovations resulting from giant virus association could involve novel metabolic functions. Especially in GVMAGs, a vast diversity of metabolic genes has been found (Schulz 2020, Moniruzzaman 2020) including many involved in carbon and nitrogen metabolism. Assessing the genetic and functional impact of giant virus infection on fungal metabolism should be a future research priority. To understand the impact of the viruses, we need to resolve when and under what conditions the viruses are expressed during the life cycle.

Notably, the most contiguous viral genomes were found in *Allomyces* spp., from cultures that were recovered from filter paper stocks after 30 years of storage. This fact demonstrates desirable aspects of this fungus that bolster motivation for its revival as a model system. *Allomyces* has historically, and currently though with much less frequency, been a model for studying biological phenomena such as anisogamy (Olsen 1984; Phadke 2018), sensorimotor systems including phototaxis and chemotaxis (Machlis 1969; Swafford and Oakley 2018), and even cellular differentiation (Cantino and Lovett 1964). The development of *Allomyces* as a model for studying giant viruses has enormous potential. Much of what is known about giant virus biology has been learned from protist-virus interactions or metagenomics where the host is unknown. The development of additional laboratory model systems of giant viruses and their natural hosts could advance our understanding of virus-host interactions, including immune response, viral “reprogramming” of host cells, metabolic impacts, and their ecosystem-level consequences.

The discovery of giant viruses in fungi that are restricted primarily to the zoosporic fungi inspire numerous questions that beg further investigation. How is viral expression and virion production related to the host life cycle, and vice-versa? What are the metabolic impacts of infection, and how important are they on the global scale? What defense systems do fungi employ against giant viruses, and do they explain why Dikarya fungi are not infected? Giant virus model systems in zoosporic fungi may be the key to exposing these latent revelations.

## Data Availability

*Mycodnaviridae* genome assemblies and alignment for the *Nucleocytoviricota* phylogeny were deposited in Deep Blue Data, the University of Michigan’s institutional data repository (DOI: https://doi.org/10.7302/p5ay-ng61). Fungal genome assemblies, and RNAseq data are available in NCBI BioProject PRJNA1257431.

## Acknowledgements

Cultures and DNAs were obtained from the Collection of Zoosporic Eufungi at the University of Michigan. Next-generation sequencing was performed at the Advanced Genomics Core at the University of Michigan We thank Mia Sinks for technical assistance with *Allomyces* cultures. JMM thanks Nina Wale for thoughtful conversation and invaluable writing companionship. This project was funded in part by the National Science Foundation grants DEB-1929738 and DEB-2403677 to TYJ and DEB-1557110 and EF-2125066 to JES. Some computations were performed using the computer clusters and data storage resources of the UC Riverside High Performance Computing Cluster, which is funded by grants from NSF (MRI-2215705, MRI-1429826) and NIH (1S10OD016290-01A1). KRA was supported by the following National Institutes of Health training grant: “Michigan Predoctoral Training in Genetics” [T32GM007544]. JES, TYJ, and JH are fellows and co-director (JH) of the CIFAR program Fungal Kingdom: Threats & Opportunities, and these studies were supported in part by a CIFAR Catalyst Award. The work conducted by the U.S. Department of Energy Joint Genome Institute (https://ror.org/04xm1d337), a DOE Office of Science User Facility, is supported by the Office of Science of the U.S. Department of Energy operated under Contract No. DE-AC02-05CH11231. Efforts by SS, VY, and JH were supported in part by NIH/NIAID R01 grants AI39115-27 and AI050113-20.

## Author Contributions

JES, TYJ, and JMM conceived of and planned the study. Culturing and DNA preparation was conducted by DRS, MOQ, and JMM. RNA preparation, PCR, transcriptome assembly, viral content identification, and orthology analysis was conducted by JMM. Giant virus phylogeny and distribution analyses were performed by FS. Genome assembly and annotation was conducted by JMM, KRA, and SR. Transcriptome analysis was conducted by SR and YV. Nanopore sequencing DNA extraction, as well as genome assembly and analysis of strain Burma1F was conducted by VY, SS and JH. Summary statistics were conducted by JES and JMM. Growth experiments were performed by TYJ. Manuscript was written by JMM with contributions from TYJ. All authors read and approved the final manuscript.

## Methods

### Fungal strains, nucleic acid preparation, and Illumina sequencing

We initially searched 135 genomes in this study including 131 fungi, 92 of which are zoosporic *(Table S1)*. For full details about DNA preparation and sequencing of these isolates see (Amses 2022). An additional nine strains of *Allomyces* spp. were whole-genome sequenced from DNA archived in the CZEUM collection (Simmons 2020). Library preparation and sequencing was performed by the Advanced Genomics Core at the University of Michigan by NovaSeq S4 300 cycle on 4% of a flowcell.

*Allomyces javanicus* California12 and *Allomyces arbusculus* Burma1F were grown in PmTG liquid media for 7-10 days at 22□with shaking at 180 rpm. Tissue was subsampled with sterile tweezers and immediately frozen and ground to a powder in liquid nitrogen. Approximately 50 mg was apportioned to 1.5mL tubes, 1 mL of TRIzol applied, and stored at -20□until further processing. For *A. arbusculus* Burma1F, the remaining tissue was decanted, rinsed once with sterile water, and replenished with sterile water. After overnight incubation at 22□, tissue was decanted, rinsed once with sterile water, subsampled and processed as described above, then transferred to a sterile tea strainer and submerged in 50 mL sterile water for two hours on the benchtop. The tea strainer was removed, and the remaining liquid observed under a microscope for zoospore presence, followed by centrifugation at 4000x g and 4□for 12 minutes. The zoospore pellet was decanted, 1mL of TRIzol added, ground with a pestle, then stored at -20□. The RNA extraction for all samples was completed per manufacturer protocol. Total RNAs were quality-checked on a Tapestation and had RIN values between 6.8-10. Samples were Poly-A enriched, and libraries were prepared and sequenced by the University of Michigan Advanced Genomics Core by NovaSeq S4 300 cycle on 1% of the flowcell.

### Genome and transcriptome assembly, annotation, and statistics

Transcriptome data were quality filtered by removing reads with Phred score <20 using Fastx-toolkit (Hannon 2010), then assembled *de novo* using Trinity 2.12 (Grabherr 2011). Hisat2 v.2.2.1 (Kim 2019) was used for the RNAseq analysis: the hisat2-build function was used to build an index from genome assemblies, then RNA sequencing reads were aligned to the indexed assemblies using default parameter settings. Alignments were visualized with SeqMonk v.1.48.1 (RRID:SCR_001913).

Paired-end genomic data of *Allomyces* strains were quality-filtered by trimmomatic (Bolger, Lohse, and Usadel 2014) with a sliding window of 4bp and threshold of Phred 20, then assembled by Spades 3.15.5 (Prjibelski 2020) and annotated with both Prodigal v. 2.6.3 (Hyatt 2010) and Augustus 3.2.1 (Stanke 2006) trained with *Rhizopus oryzae*.

For the genome plots of *Allomyces arbusculus* Burma 1F *and Allomyces javanicus* California12, we used Prodigal v. 2.6.3 (Hyatt 2010) for gene prediction of soft-masked viral contigs with inputs from Funannotate v. 1.8.16 (Palmer and Stajich 2023) which is a fungal gene annotation tool and GeneMarkS (http://exon.gatech.edu/genemark/index.html) using the viral option for input sequence type. GC skew was calculated in a 5-kb sliding window with a 1-kb step size. Circos v. 0.54 (Krzywinski 2009) was used to construct the circular genome visualization.

### Identification of viral content

We used the program ViralRecall (Aylward and Moniruzzaman 2021) to identify putative viral genes, viral contigs, and viral regions within contigs. We first ran ViralRecall 1.0 on all assemblies from Amses 2022 (n = 131) with both stringent (-s 15 -m 30 -v 10) and relaxed (-s 15 -m 2 -v 2) parameters–the latter for poorer quality genome assemblies–and identified isolates containing MCP homologs with e-value <1e^-10^. ViralRecall searches gene predictions generated using Prodigal (Hyatt 2010); we also performed hmmsearches (HMMER3 v3.1b2; S. Eddy 2009) with the ViralRecall 2.0 GVOG hmm on a subset of our Augustus-predicted genomes. We found no differences in the number of MCPs called, and so continued characterizations for the isolates that these initial ViralRecall results predicted at least one MCP homolog. For each MCP-containing contig, we calculated GC content, intron density per the Augustus annotation, and performed BLAST searches (Altschul 1990) on both gene prediction datasets.

To aid in distinguishing viral genomes from within the fungal assemblies, we searched the 16 prodigal annotated genomes previously identified to contain MCPs with HMMER3 (Eddy 2009) as in Schulz 2020 using hmms of a subset of the nucleocytoplasmic viral orthologous genes (NCVOGs)— the 20 NCVOGs most likely to have been vertically inherited (Yutin 2009). We made individual gene trees including all hits with e-values less than 1e-10 by aligning gene sequences with MAFFT version 7 using the E-INS-i algorithm (Katoh and Standley 2013), trimming the resulting alignments with the -automated1 method in TrimAl (Capella-Gutiérrez 2009), and reconstructing trees with the approximately maximum-likelihood approach implemented in FastTree (Price, Dehal, and Arkin 2009) with 100 bootstrap replicates, rooting at the midpoint. We also ran the latest version of ViralRecall (2.0; Aylward and Moniruzzaman 2021) on these 16 assemblies and used these results in combination with genome statistics to discern putatively viral contigs, which we classified as belonging to a viral genome if they met the following criteria: ViralRecall score >2, viral region length (per ViralRecall) >70% of the total contig length, and intron density > 25% different from the host average. One isolate appeared to contain two distinct NCLDVs; we used MetaBAT2 (Kang 2019) on the entire genome of *Allomyces javanicus* California12 under the default settings to separate the two NCLDV genomes.

We conducted BLASTP searches of the hybrid proteomes against nr to identify homologs (e-value <1e-10). Because many symbiont genes are likely misidentified as their hosts’ in the databases, we labeled a gene as “viral” or “bacterial” if any of the top hits were to a virus or bacterium. Some genes have homologs only in the fungi which we have shown also encode MCP; we labeled these “putative FNCLDV”.

To assess prevalence of fungal giant virus-like MCP throughout the fungal kingdom, we used blastp to search all unmasked fungal genomes in Mycocosm (Grigoriev 2014) using MCP of *Allomyces arbusculus* Burma1F as query (total 2,543 genomes; conducted on Dec 30, 2023). We also searched these Mycocosm genomes using the following protein models: PF04451 (Large eukaryotic DNA virus major capsid protein), IPR007542 (Major capsid protein, C-terminal), IPR031654 (Major capsid protein, N-terminal), and IPR016112 (Group II dsDNA virus coat/capsid protein) (conducted on Jan 13, 2026). The first three profile searches yielded hits only to zoosporic fungi, with the exception of a gene in *Dimargaris cristalligena* (hit to IPR007542). Genes with homology to IPR016112 were found across fungal lineages, including in Dikarya. These genes hits, if in zoosporic fungi, typically shared annotations with PF04451 and/or IPR007542; hits in non-zoosporic lineages never shared annotations with other viral or structural proteins, suggesting non-viral function. Interestingly, searching with the Major Capsid protein N-terminal (IPR031654) yielded no results, suggesting that fungal MCPs have a unique N-terminus.

### Giant virus phylogeny, orthology analysis, and KilA-N domain search

To build the giant virus species tree the nsgtree pipeline was used (https://github.com/NeLLi-team/nsgtree) on genome representatives of the viral phylum Nucleocytoviricota (Aylward 2021). In brief, 7 giant virus orthologous groups (GVOGs; Aylward 2021) were identified using hmmsearch, extracted GVOGs were aligned with MAFFT (version 7.31; Katoh and Standley 2013), trimmed with trimal (-gt 0.1, v1.4; Capella-Gutiérrez, Silla-Martínez, and Gabaldón 2009) and concatenated. To limit the amount of missing data we removed genomes that had less than 4 out of 7 GVOGs. A species tree was built from the supermatrix alignment using IQ-Tree (version 2.03); (Minh 2020; Kalyaanamoorthy 2017) with LG+I+F+G4 and ultrafast bootstrap replications (Hoang 2018) and visualized in iToL (Letunic and Bork 2021).

To analyze the global distribution of *Mycodnaviridae* we used the MCP as a query for a blastp using diamond (--query-cover 30 --subject-cover 30; Buchfink, Xie, and Huson 2015) against all MCP that we previously extracted (Schulz 2020) from publicly available IMG metagenomes (Chen 2023). All blastp hits were then extracted and used as query for another blastp against a database consisting of all MCP extracted from representative giant virus genomes (Aylward 2021) and the initial *Mycodnaviridae* MCPs. We considered as bidirectional best blastp hits (BBH) the environmental MCP that had the best hit based on bitscore to the initial *Mycodnaviridae* MCPs. Geolocation of the BBHs was then used to plot the distribution of MCPs related to *Mycodnaviridae* on a world map projection.

To search for homologs of the viral KilA-N domain in *Mycodnaviridae* genomes, the source data from figure 6 of Medina 2016 (219 protein sequences) was downloaded. Sequences were aligned using mafft (version 7.31; Katoh and Standley 2013), and an hmmprofile made with HMMER hmmbuild (Eddy 2009), which was then used to hmmsearch the fungal giant virus genomes.

Orthofinder 2.5.2 (Emms and Kelly 2019) was used to identify orthologous gene clusters from the prodigal predictions of the most complete viral genomes, determined as the viral genomes containing the greatest number (at least nine) of the NCVOGs20. Visualization of the shared orthologous gene clusters was by adaptation of https://commons.wikimedia.org/wiki/File:Symmetrical_5-set_Venn_diagram.svg. Sequences of orthologous gene clusters were fed into EggNOG-mapper 5.0 (Huerta-Cepas 2019) with default parameters but enabling SMART annotations.

### *Allomyces* capsid PCR and growth experiment

Primers were designed from an alignment of the MCP sequences of *A. javanicus* California12 and *A. arbusculus* Burma1F (F 5’ TACACCAACTTTGCGATGGA3’ R 5’ GTAGCTCGTTGACCCGACAT3’). DNA was provided by the Collection of Zoosporic Eufungi at the University of Michigan. A standard PCR was performed using GoTaq (Promega) and the products visualized by agarose gel electrophoresis for absence/presence. Burma1F was used as a positive control, and Burma3-35 served as a negative control based on absence of the gene in the sequenced genome.

We chose 6 strains that tested positive for MCP (BEA2, Burma1F, Cali8, Cali12, FijiF1, NorthCaro2), and 6 testing negative (ATCC10983, Burma3-35, CubaS7, DJ02, DJ07, Mex58) to determine phenotypic effects of the gene’s presence. Strains were grown in triplicate on Emerson’s YpSS media (Emerson 1958). The inoculum for each plate was made by growing strains for 48 hours on fresh YpSS and removing a small circular inoculum by punching discs with the back of a glass pipette. After 72 hrs of growth at 35 C, linear growth was measured for each plate with two perpendicular radii. We also estimated biomass accumulation of the same strains using liquid YpSS. Triplicate 250 ml flasks with 100 ml of media were inoculated with 4 discs cut using the back of glass pipettes. The strains were grown at 35 C for 48 hours with shaking at 120 rpm. The mycelium was harvested using miracloth filtration, transferred to pre-weighed filter papers, dried completely, and dry weight was measured. A final experiment measured thermal maxima of strains by inoculation of YpSS plates and placing them at multiple temperatures: 38, 40, 42, 44, and 46 C. Duplicate plates were prepared from 9 day old YpSS plates using discs excised using glass pipettes. Plates were scored for evidence of growth, where < 2 mm of growth after 1 week was considered negative.

### *Allomyces* qPCR and transmission experiments

The starting material for these experiments were resting sporangia that had been dried down onto filter paper and stored for at least 30 years when they were provided to the Fungal Genetics Stock Center by Lene Lange. Single meiospore isolates of Cali12, Burma1F, and Cali8 were obtained by placing the resting sporangia into sterile deionized water. After a period of 1-4 days, meiospores were released and plated onto YpSS medium supplemented with penicillin and streptomycin sulfate (200 mg/L). Individual germlings were found under a stereoscope and subcultured.

We estimated the amount of viral DNA in these samples by qPCR by comparison of the copy number of the MCP to actin genes in DNA extracts. Samples for qPCR were grown in ¼ strength liquid YpSS for 6-12 days at either 22 C or 37 C. DNA was extracted using a modified CTAB protocol, and extracts were quantified using a qubit fluorimeter and diluted to 0.1 ng/ul.Primers and probes were designed using PrimerQuest (IDT) (Supplementary Table 2). Probes were IDT PrimeTime probes with double quenchers. To create a standard curve we used synthetic DNA fragments (gblocks) of a known concentration. Reactions were conducted using RADIANT Lo-ROX 2x master mix (Alkali Scientific) on a QuantStudio 3 Thermal Cycler (Applied Biosystems, Thermo Fisher Scientific).

RNA was extracted from liquid and plate YPSS/4 cultures using TRIzol per manufacturer recommendations on tissue that was first ground to a powder in liquid nitrogen, then treated with DNase. cDNA was generated using LunaScript RT SuperMix Kit (NEB) with random hexamers.

We performed an experiment to test whether we could transmit the virus from Cali8 to a virus-free *Allomyces* strain (Mex58). We grew two subcultures of Cali8 that had been shown to have high viral loads (m1 and m1-3) and Burma3-35 (virus-free) in liquid Y/4 and grown for 13 days. We removed the mycelium from the media and grinded the mycelium in an eppendorf tube with a micropestle until it was homogenous. Both the homogenate and the medium were then filtered using a 0.2 um filter and these filtrates applied to pools of Mex58 gametes which had been released from a gametophyte culture by flooding with water. We combined 0.5 ml of gametes (> 10^3^) and either 0.5 ml of filtered supernatant or 0.15 ml of filtered homogenate plus 1 ml of fresh YpSS in a multiwell plate and incubated the plate for 4 days. After this time, the cells were transferred to 1 ml fresh media with a 1/10 dilution or a 1/100 dilution and grown at 22 C for 16 days at which point a sporophyte mycelium filled the wells. There were 12 total experiments: 2 dilutions (1/10 or 1/100, 3 strains (m1, m1-3, and Burma3-35), and two inoculum types (homogenized cells or supernatant). The total mycelium of the 12 experiments was harvested, DNA extracted using a modified CTAB method, and qPCR of the MCP and actin genes used to assess whether successful transmission occurred.

### Nanopore Sequencing and Genome Assembly

To obtain High Molecular Weight DNA for long read sequencing, the strain was first grown on YpSS solid medium for 3 weeks. A block (3mm x 3mm) from the edge of the growing culture was then excised and immersed in DS buffer to induce zoospore release. The spore suspension was used to inoculate 100 ml fresh YpSS liquid medium, which was then incubated for 36 hours at 33 °C in a floor shaker. After incubation, the cells were collected with strainer, washed once with PBS, frozen at -80 °C, and lyophilized. The frozen-dry cells were broken into powder using 4 mm glass beads and vortex at the maximum speed. Genomic DNA was then extracted using a modified CTAB protocol, and DNA quality was inspected with CHEF gel electrophoresis as previously described (https://doi.org/10.1073/pnas.2416656121).

Purified DNA was processed for library preparation and sequencing on an Oxford Nanopore MinION Mk1C device using R10 flow cells (FLO-MIN114) with MinKNOW UI (v23.07.12) following the sequencing kit guidelines. The generated Pod5 data was subjected to base calling using Dorado (v0.6.0) followed by genome assembly using Canu (v2.1.1). The resulting genome assembly was analyzed for the presence of the capsid sequence using BLASTn analysis and transposons were identified using Repeatmasker (v4.0.7; http://www.repeatmasker.org), supplemented with Dfam (v3.3), RepBaseRepeatMaskerEdition-20181026 libraries, and RepBase EMBL database (v26.04; Bao 2015, Storer 2021). CpG methylation analysis was performed using the nanopore Modkit tool (v0.2.5-rc1).

