## Supporting Information for "Discovery of *Mycodnaviridae*, a clade of giant viruses that persistently infect zoosporic fungi"

### Supplemental Tables and Figures

Figure S1. Phylogeny spotlighting the zoosporic fungi, adapted from Amses *et al.* 2022. Red stars indicate species found to have integrated Major Capsid Proteins.

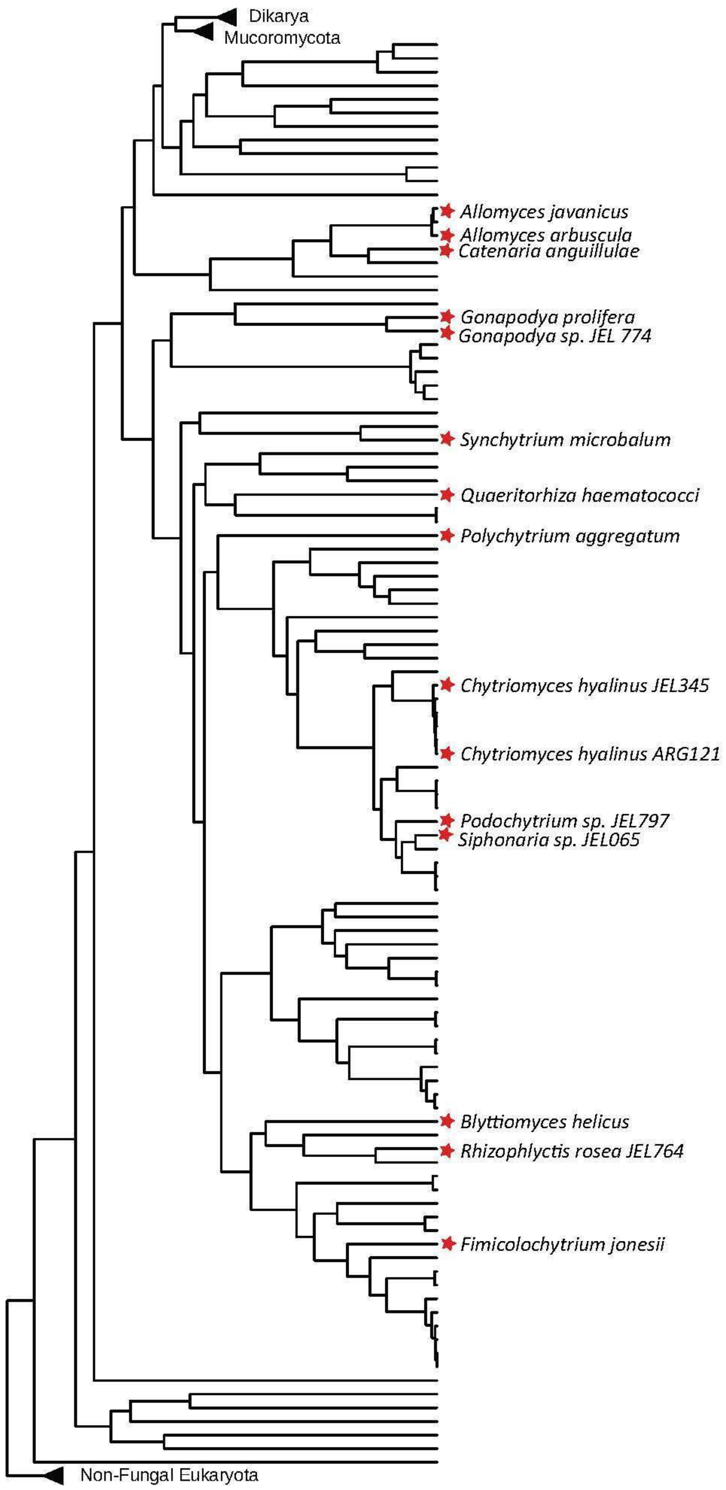

Table S1: Shown is the number of ten key giant virus orthologous genes (GVOGs) found across fungal genomes by HMMsearch (eval < 1e-10).

| Isolate | Phylum | MCP | VLTF3 | D5 | A32 | SFII | mRNAc | RNR | RNAPS | RNAPL | PolB |
|---|---|---|---|---|---|---|---|---|---|---|---|
| Arthrobotrys_oligospora_ATCC_24927 | Ascomycota | 0 | 0 | 0 | 0 | 0 | 0 | 0 | 8 | 8 | 4 |
| Candida_arabinofermentans_NRRL_YB-2248 | | 0 | 0 | 0 | 0 | 2 | 2 | 1 | 3 | 3 | 4 |
| Hortaea_werneckii_EXF-2000 | | 0 | 0 | 0 | 0 | 4 | 4 | 2 | 6 | 10 | 10 |
| Neurospora_crassa_OR74A | | 0 | 0 | 0 | 0 | 1 | 2 | 1 | 2 | 4 | 6 |
| Pecoramyces_ruminatium_C1A | | 0 | 0 | 4 | 0 | 0 | 1 | 0 | 11 | 7 | 5 |
| Saccharomyces_cerevisiae_S288C | | 0 | 0 | 0 | 0 | 2 | 2 | 2 | 3 | 3 | 4 |
| Saccharomycopsis_capsularis_NRRL_Y-17638 | | 0 | 0 | 0 | 0 | 2 | 2 | 2 | 3 | 3 | 4 |
| Schizosaccharomyces_pombe_972h | | 0 | 0 | 0 | 0 | 1 | 2 | 2 | 3 | 3 | 4 |
| Yarrow_lipolytica_CLIB122 | | 0 | 0 | 0 | 0 | 2 | 2 | 1 | 3 | 3 | 4 |
| Armillaria_ostoyae_C18_9 | Basidiomycota | 0 | 0 | 0 | 0 | 0 | 0 | 23 | 8 | 7 | 2 |
| Coprinopsis_cinerea_okayama7_130 | | 0 | 0 | 0 | 0 | 0 | 1 | 2 | 4 | 3 | 3 |
| Mixia_osmundae | | 0 | 0 | 0 | 0 | 0 | 0 | 4 | 4 | 12 | 4 |
| Puccinia_graminis_f._sp._tritici_CRL_75-36-700-3 | | 0 | 0 | 0 | 0 | 0 | 0 | 0 | 7 | 9 | 3 |
| Ustilago_maydis_521 | | 0 | 0 | 0 | 0 | 2 | 2 | 1 | 4 | 3 | 4 |
| Backusella_circina_FSU_941 | Mucoromycota | 0 | 0 | 0 | 0 | 1 | 0 | 3 | 7 | 8 | 5 |
| Bifiguratus_adelaidae_AZ0501 | | 0 | 0 | 1 | 0 | 0 | 1 | 1 | 11 | 8 | 7 |
| Gigaspora_rosea_DAOM_194757 | | 0 | 0 | 0 | 0 | 0 | 0 | 3 | 7 | 7 | 77 |
| Hesseltinella_vesiculosa_NRRL_3301 | | 0 | 0 | 0 | 0 | 1 | 0 | 1 | 8 | 8 | 3 |
| Lichtheimia_corymbifera_FSU_9682 | | 0 | 0 | 0 | 0 | 1 | 0 | 4 | 7 | 10 | 2 |
| Lobosporangium_transversale_NRRL_3116 | | 0 | 0 | 0 | 0 | 1 | 0 | 2 | 12 | 10 | 6 |
| Mortierella_elongata_AG-77 | | 0 | 0 | 1 | 0 | 1 | 1 | 3 | 10 | 9 | 7 |
| Mucor_circinelloides_f._circinelloides_1006PhL | | 0 | 0 | 0 | 0 | 1 | 0 | 3 | 7 | 5 | 6 |
| Phycomyces_blakesleeanus_NRRL_1555 | | 0 | 0 | 0 | 0 | 1 | 0 | 1 | 6 | 9 | 5 |
| Rhizophagus_irregularis_DAOM_181602 | | 0 | 0 | 0 | 0 | 0 | 0 | 2 | 12 | 9 | 44 |
| Rhizopus_delemar_RA_99-880 | | 0 | 0 | 0 | 0 | 1 | 0 | 2 | 14 | 11 | 5 |
| Rhizopus_microsporus_var._microsporus_ATCC_52814 | | 0 | 0 | 0 | 0 | 1 | 0 | 1 | 5 | 8 | 4 |
| Saksenaea_vasiformis_B4078.G233 | | 0 | 0 | 0 | 0 | 1 | 0 | 2 | 7 | 8 | 5 |
| Syncephalastrum_racemosum_NRRL_2496 | | 0 | 0 | 0 | 0 | 1 | 1 | 2 | 11 | 8 | 5 |
| Umbelopsis_ramanniana_AG | | 0 | 0 | 0 | 0 | 1 | 0 | 2 | 8 | 9 | 4 |
| Basidiobolus_heterosporus_B8920.N168 | Zoopagomycota | 0 | 0 | 3 | 0 | 0 | 0 | 3 | 12 | 10 | 5 |
| Basidiobolus_meristosporus_CBS_931.73 | | 0 | 0 | 0 | 0 | 1 | 0 | 3 | 9 | 11 | 4 |
| Coemansia_reversa_NRRL_1564 | | 0 | 0 | 0 | 0 | 2 | 2 | 1 | 4 | 6 | 4 |
| Conidiobolus_coronatus_NRRL_28638 | | 0 | 0 | 1 | 0 | 0 | 0 | 1 | 5 | 4 | 6 |
| Conidiobolus_thromboides_FSU_785 | | 0 | 0 | 2 | 0 | 1 | 2 | 1 | 9 | 7 | 3 |
| Linderina_pennispora_ATCC_12442 | | 0 | 0 | 0 | 0 | 3 | 2 | 1 | 9 | 10 | 6 |
| Martensiomyces_pterosporus_CBS_209.56 | | 0 | 0 | 0 | 0 | 2 | 1 | 1 | 3 | 4 | 4 |
| Piptocephalis_cylindrospora_RSA_2659 | | 0 | 0 | 0 | 0 | 0 | 1 | 0 | 0 | 3 | 1 |
| Ramicandelaber_brevisporus_CBS_109374 | | 0 | 0 | 0 | 0 | 2 | 2 | 1 | 5 | 5 | 5 |
| Syncephalis_fuscata_S228 | | 0 | 0 | 0 | 0 | 0 | 0 | 1 | 11 | 5 | 0 |
| **Allomyces_arbuscula_Burma_1F** | Blastocladiomycota | **2** | **1** | **2** | **2** | **1** | **1** | **2** | **4** | **11** | **6** |
| **Allomyces_javanicus_California_12** | | **5** | **2** | **9** | **4** | **3** | **2** | **4** | **4** | **11** | **9** |
| Allomyces_macrogynus_ATCC_38327 | | 0 | 0 | 2 | 9 | 0 | 0 | 2 | 7 | 14 | 9 |
| Blastocladiella_britannica | | 0 | 0 | 0 | 0 | 0 | 0 | 2 | 4 | 6 | 3 |
| **Catenaria_anguillulae_PL171_v2.0** | | **1** | **0** | **1** | **0** | **1** | **1** | **5** | **8** | **12** | **8** |
| Coelomomyces_lativittatus_CIRM-AVA-1-Meiospore | | 0 | 0 | 0 | 0 | 0 | 0 | 2 | 5 | 4 | 4 |
| Paraphysoderma_sedebokerense_JEL0847 | | 0 | 0 | 5 | 3 | 1 | 2 | 3 | 10 | 12 | 9 |
| Anaeromyces_robustus | | 0 | 0 | 4 | 0 | 0 | 4 | 0 | 8 | 8 | 6 |
| Batrachochytrium_dendrobatidis_JAM81 | | 0 | 0 | 0 | 0 | 0 | 0 | 2 | 5 | 6 | 2 |
| Batrachochytrium_dendrobatidis_JEL423 | | 0 | 0 | 0 | 0 | 0 | 0 | 2 | 5 | 6 | 2 |
| Batrachochytrium_salamandrivorans | | 0 | 0 | 0 | 0 | 0 | 0 | 2 | 6 | 5 | 2 |
| **Blyttiomyces_helicus_single-cell_v1.0** | | **8** | **2** | **29** | **2** | **2** | **3** | **7** | **2** | **16** | **9** |
| Blyttiomyces_sp._JEL0837 | | 0 | 0 | 0 | 0 | 0 | 0 | 0 | 0 | 0 | 0 |
| Boothiomyces_macroporosum_PLAUS21 | | 0 | 0 | 1 | 0 | 0 | 0 | 3 | 10 | 21 | 6 |
| Borealophylctis_nickersoniae_WJD170 | | 0 | 0 | 36 | 1 | 0 | 0 | 1 | 11 | 6 | 2 |
| Caulochytrium_protostelioides_ATCC_52028 | | 0 | 0 | 0 | 0 | 2 | 2 | 1 | 3 | 3 | 4 |
| Chytridium_lagenaria_Arg66 | | 0 | 0 | 0 | 0 | 0 | 0 | 1 | 9 | 6 | 0 |

| Isolate | Phylum | MCP | VLTF3 | D5 | A32 | SFII | mRNAc | RNR | RNAPS | RNAPL | PolB |
|---|---|---|---|---|---|---|---|---|---|---|---|
| Chytriomyces_confervae_CBS_675.73 | Chytridiomycota | 0 | 0 | 2 | 0 | 0 | 1 | 1 | 13 | 11 | 1 |
| Chytriomyces_hyalinus_ARG085 | | 0 | 0 | 0 | 0 | 0 | 0 | 1 | 13 | 9 | 1 |
| **Chytriomyces_hyalinus_ARG121** | | **1** | **0** | **5** | **0** | **0** | **0** | **2** | **8** | **12** | **1** |
| Chytriomyces_hyalinus_JEL0117 | | 0 | 0 | 5 | 0 | 0 | 0 | 3 | 14 | 14 | 0 |
| Chytriomyces_hyalinus_JEL0176 | | 0 | 0 | 3 | 0 | 0 | 1 | 2 | 14 | 9 | 1 |
| **Chytriomyces_hyalinus_JEL0345** | | **2** | **1** | **46** | **3** | **1** | **2** | **4** | **8** | **17** | **5** |
| Chytriomyces_sp._MP_71 | | 0 | 0 | 0 | 0 | 0 | 0 | 2 | 15 | 11 | 5 |
| Cladochytrium_replicatum_JEL714 | | 0 | 0 | 0 | 0 | 0 | 1 | 2 | 12 | 11 | 1 |
| Cladochytrium_tenue_CCIBt4013 | | 0 | 0 | 0 | 0 | 0 | 0 | 4 | 13 | 13 | 2 |
| Clydaea_vesicula_JEL0476 | | 0 | 0 | 5 | 0 | 9 | 2 | 2 | 11 | 8 | 5 |
| Dinochytrium_kinnereticum_KLL_TL_06062013 | | 0 | 0 | 0 | 0 | 0 | 0 | 1 | 4 | 9 | 1 |
| Entophlyctis_helioformis_JEL805 | | 0 | 0 | 0 | 0 | 0 | 0 | 1 | 16 | 10 | 4 |
| Entophlyctis_luteolus_JEL0129 | | 0 | 0 | 1 | 0 | 0 | 0 | 4 | 10 | 15 | 2 |
| Entophlyctis_sp._JEL0112 | | 0 | 0 | 1 | 0 | 0 | 0 | 4 | 12 | 15 | 2 |
| **Fimicolochytrium_jonesii_JEL569** | | **1** | **0** | **23** | **0** | **3** | **2** | **3** | **11** | **6** | **3** |
| **Gaertneriomyces_semiglobifer_Barr_43** | | **0** | **0** | **19** | **1** | **2** | **1** | **4** | **9** | **6** | **4** |
| Gaertneriomyces_sp._JEL0708 | | 0 | 0 | 4 | 0 | 2 | 2 | 6 | 10 | 8 | 7 |
| Geranomyces_michiganensis_JEL0563 | | 0 | 0 | 4 | 0 | 1 | 1 | 2 | 8 | 10 | 5 |
| Geranomyces_variabilis_JEL0379 | | 0 | 0 | 4 | 0 | 1 | 2 | 2 | 9 | 10 | 5 |
| Geranomyces_variabilis_JEL0389 | | 0 | 0 | 7 | 0 | 1 | 1 | 1 | 8 | 9 | 4 |
| Geranomyces_variabilis_JEL0566 | | 0 | 0 | 5 | 0 | 1 | 1 | 1 | 8 | 9 | 4 |
| Geranomyces_variabilis_JEL0567 | | 0 | 0 | 4 | 0 | 1 | 1 | 2 | 9 | 12 | 4 |
| Geranomyces_variabilis_JEL559 | | 0 | 0 | 4 | 0 | 1 | 1 | 2 | 8 | 12 | 4 |
| Globomyces_pollinis-pini_Arg68 | | 0 | 0 | 0 | 0 | 0 | 0 | 1 | 9 | 9 | 1 |
| **Gonapodya_prolifera** | | **2** | **4** | **7** | **1** | **1** | **1** | **0** | **7** | **13** | **5** |
| **Gonapodya_sp._JEL0774** | | **5** | **1** | **28** | **1** | **1** | **1** | **2** | **7** | **16** | **3** |
| Gorgonomyces_haynaldii_MP57 | | 0 | 0 | 0 | 0 | 1 | 1 | 2 | 8 | 6 | 5 |
| Homolaphlyctis_polyrhiza_JEL142 | | 0 | 0 | 0 | 0 | 0 | 1 | 1 | 9 | 8 | 4 |
| Hyaloraphidium_curvatum_SAG235-1 | | 0 | 0 | 6 | 2 | 0 | 0 | 1 | 6 | 9 | 4 |
| Irineochytrium_annulatum_JEL0729 | | 0 | 0 | 0 | 0 | 3 | 0 | 5 | 10 | 9 | 4 |
| Kappamyces_sp._JEL0680 | | 0 | 0 | 0 | 0 | 0 | 2 | 2 | 8 | 9 | 5 |
| Lobulomyces_angularis_JEL0522 | | 0 | 0 | 2 | 0 | 0 | 1 | 2 | 10 | 9 | 4 |
| Lobulomyces_sp._JEL0476 | | 0 | 0 | 5 | 0 | 0 | 1 | 2 | 11 | 8 | 6 |
| Neocallimastix_californiae_G1 | | 0 | 0 | 23 | 0 | 0 | 13 | 0 | 46 | 70 | 35 |
| Nowakowskiella_sp._JEL0407 | | 0 | 0 | 0 | 0 | 0 | 0 | 2 | 26 | 10 | 0 |
| Obelidium_mucronatum_JEL802 | | 0 | 0 | 0 | 0 | 0 | 0 | 4 | 8 | 18 | 1 |
| Olpidium_bornovanus_UCB_F19785 | | 0 | 0 | 1 | 0 | 0 | 1 | 1 | 15 | 16 | 4 |
| Olpidium_sp._PSC023 | | 0 | 0 | 1 | 0 | 2 | 0 | 1 | 5 | 4 | 3 |
| Paranamyces_uniporus_JEL0695 | | 0 | 0 | 0 | 0 | 1 | 1 | 3 | 19 | 17 | 7 |
| Phlyctochytrium_bullatum_JEL0754 | | 0 | 0 | 1 | 0 | 0 | 0 | 2 | 11 | 10 | 1 |
| Phlyctochytrium_planicorne_JEL0388 | | 0 | 0 | 0 | 0 | 0 | 0 | 0 | 9 | 8 | 1 |
| Physocladia_obscura_JEL0513 | | 0 | 0 | 6 | 0 | 0 | 1 | 2 | 8 | 10 | 1 |
| Piromyces_finnis | | 0 | 0 | 7 | 1 | 0 | 3 | 0 | 16 | 22 | 7 |
| Piromyces_sp._E2 | | 0 | 0 | 2 | 0 | 0 | 0 | 0 | 9 | 10 | 7 |
| **Podochytrium_sp._JEL0797** | | **1** | **1** | **1** | **0** | **0** | **0** | **2** | **6** | **9** | **1** |
| **Polychytrium_aggregatum_JEL109** | | **3** | **1** | **27** | **1** | **0** | **0** | **9** | **14** | **14** | **4** |
| Powellomyces_hirtus_BR81 | | 0 | 0 | 4 | 0 | 1 | 1 | 5 | 8 | 8 | 4 |
| Powellomyces_hirtus_CBS809.83 | | 0 | 0 | 12 | 0 | 2 | 1 | 0 | 9 | 7 | 4 |
| **Quaeritorhiza_haematococci_JEL0916** | | **8** | **1** | **85** | **0** | **2** | **2** | **3** | **12** | **9** | **3** |
| Rhizoclosmatium_globosum_JEL800 | | 0 | 0 | 6 | 0 | 0 | 0 | 2 | 13 | 22 | 1 |
| Rhizoclosmatium_hyalinum_JEL0917 | | 0 | 0 | 2 | 0 | 0 | 0 | 2 | 12 | 15 | 3 |
| Rhizophlyctis_rosea_JEL0318 | | 0 | 0 | 35 | 1 | 0 | 0 | 4 | 12 | 6 | 1 |
| **Rhizophlyctis_rosea_JEL0764** | | **1** | **0** | **27** | **2** | **1** | **0** | **3** | **10** | **7** | **1** |
| Rhizophydium_sp._JEL0728 | | 0 | 0 | 0 | 2 | 0 | 0 | 5 | 15 | 23 | 6 |
| Rhizophydium_sp._JEL0801 | | 0 | 0 | 0 | 0 | 0 | 0 | 1 | 12 | 11 | 3 |
| Rhizophydium_sp._JEL0829 | | 0 | 0 | 1 | 0 | 0 | 2 | 2 | 11 | 11 | 6 |

| Isolate | Phylum | MCP | VLTF3 | D5 | A32 | SFII | mRNAc | RNR | RNAPS | RNAPL | PolB |
|---|---|---|---|---|---|---|---|---|---|---|---|
| Rhizophydium_sp._JEL0838 | | 0 | 0 | 2 | 5 | 1 | 0 | 3 | 7 | 11 | 6 |
| Rhizophydium_sp._JEL0842 | | 0 | 0 | 0 | 0 | 0 | 0 | 2 | 6 | 9 | 1 |
| Rhizophydium_sp._JEL0866 | | 0 | 0 | 6 | 3 | 1 | 3 | 3 | 9 | 13 | 9 |
| **Siphonaria_sp._JEL0065** | | **3** | **0** | **5** | **0** | **0** | **1** | **3** | **20** | **25** | **7** |
| Spizellomyces_palustris_CBS455.65 | | 0 | 0 | 2 | 0 | 1 | 2 | 3 | 8 | 7 | 4 |
| Spizellomyces_palustris_CBS455.65 | | 0 | 0 | 2 | 0 | 1 | 2 | 3 | 8 | 7 | 4 |
| Spizellomyces_punctatus_DAOM_BR117 | | 0 | 0 | 5 | 0 | 1 | 1 | 2 | 10 | 8 | 4 |
| Synchytrium_endobioticum_MB42 | | 0 | 0 | 0 | 0 | 0 | 0 | 1 | 7 | 7 | 2 |
| **Synchytrium_microbalum_JEL0517** | | **3** | **1** | **13** | **1** | **1** | **1** | **3** | **6** | **4** | **1** |
| Thoreauomyces_humboldtii_JEL0095 | | 0 | 0 | 22 | 1 | 1 | 1 | 2 | 9 | 6 | 3 |
| Triparticalcar_arcticum_BR59 | | 0 | 0 | 37 | 0 | 1 | 1 | 2 | 4 | 5 | 3 |
| Unknown_unknown_JEL0888 | | 0 | 0 | 0 | 0 | 0 | 0 | 2 | 4 | 6 | 5 |
| Zopfochytrium_polystomum_WB228 | | 0 | 1 | 45 | 0 | 0 | 1 | 2 | 9 | 11 | 6 |
| Paramicrosporidium_saccamoebae_KSL3 | Cryptomycota | 0 | 0 | 0 | 0 | 2 | 2 | 1 | 3 | 4 | 3 |
| Rozella_allomycis_CSF55 | | 0 | 0 | 0 | 0 | 1 | 0 | 1 | 8 | 9 | 4 |
| Rozella_multimorpha | | 0 | 0 | 0 | 0 | 0 | 1 | 1 | 10 | 10 | 4 |
| Rozella_sp._PSC023 | | 0 | 0 | 0 | 0 | 1 | 0 | 2 | 4 | 5 | 6 |
| Mitosporidium_daphniae_UGP3 | | 0 | 0 | 0 | 0 | 0 | 0 | 1 | 7 | 10 | 5 |
| Fonticula_alba_ATCC_38817 | Outgroup | 0 | 0 | 0 | 0 | 0 | 1 | 1 | 7 | 11 | 4 |
| Capsaspora_owczarzaki_ATCC_30864 | | 0 | 0 | 3 | 0 | 2 | 2 | 3 | 2 | 4 | 5 |
| Drosophila_melanogaster | | 0 | 0 | 0 | 0 | 0 | 1 | 1 | 4 | 9 | 2 |
| Paraphelidium_tribonemae_X-108 | | 0 | 0 | 5 | 1 | 11 | 3 | 11 | 25 | 35 | 7 |
| Monosiga_brevicolis_MX1 | | 0 | 0 | 0 | 1 | 1 | 0 | 1 | 5 | 5 | 3 |

Fig S2: Zoosporic fungi found with Major Capsid Proteins (MCPs) demonstrated variation in genomic properties of the viral introgression. The contig containing the MCP in *Chytriomyces hyalinus* JEL345 (lower left) is a mosaic of bacterial, viral, and fungal genes, with reduced GC content and intron density compared with fungal nuclear contigs from the same genome assembly (top left). Contrarily, the MCP-containing contig in *Podochytrium* sp. JEL797 (lower right) is composed of predominantly fungal genes, with similar GC content and intron density as a fungal control contig from the same assembly (upper right).

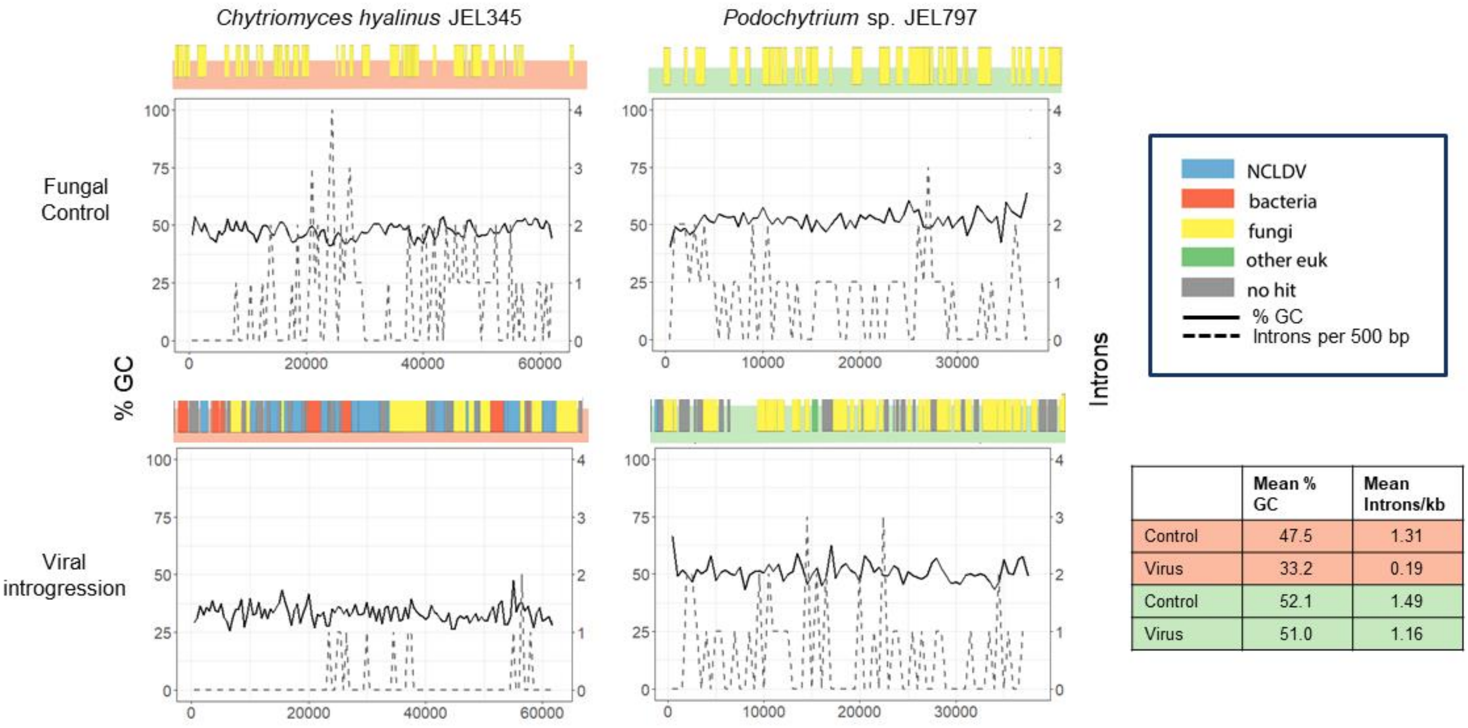


| | Mean % GC | Mean Introns/kb |
|---|---|---|
| Control | 47.5 | 1.31 |
| Virus | 33.2 | 0.19 |
| Control | 52.1 | 1.49 |
| Virus | 51.0 | 1.16 |

Fig S3: Circos plot of *Allomyces javanicus* California12 mycodnavirus 1, created as for *A. arbusculus* Burma1F (see methods).

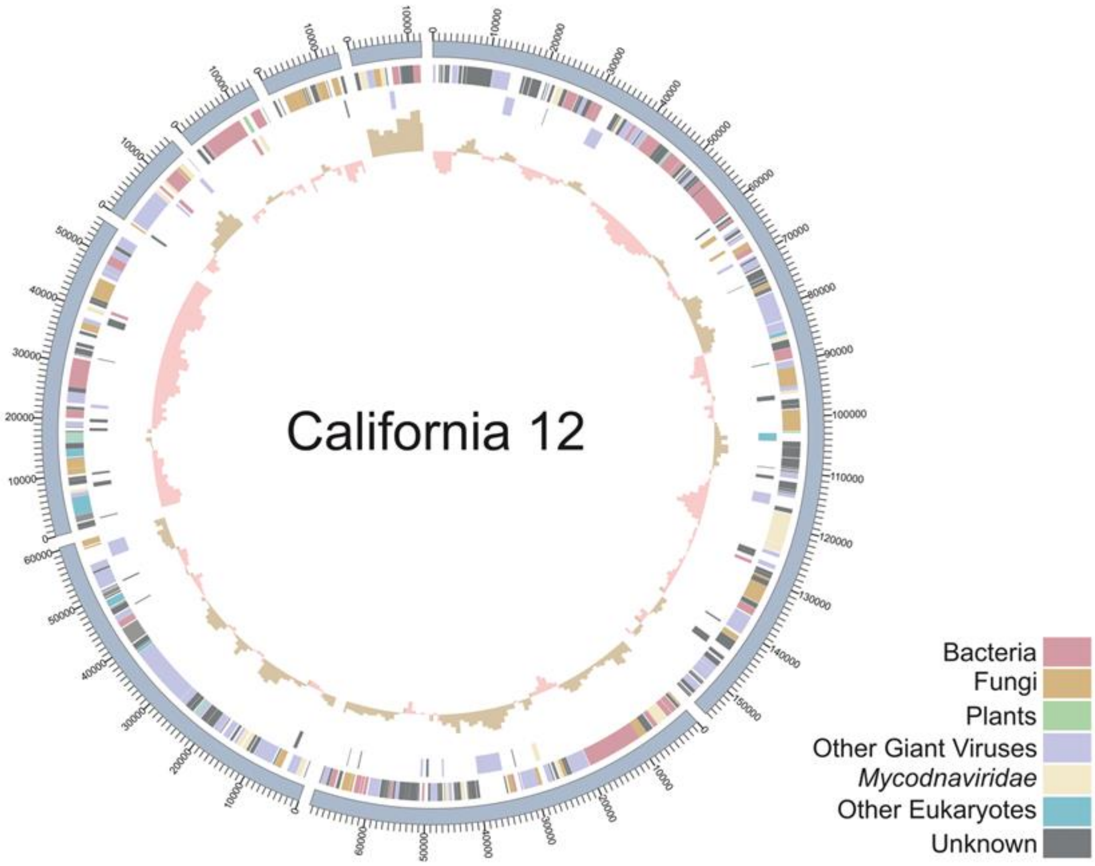

Figure S4. Left panel: Percent GC content by Coverage plots of *A. arbusculus* BEA2 (upper) and *A. arbusculus* Cali8 Illumina assemblies (lower). Contigs containing MCP genes are shown in blue, each point is sized according to its length. The contigs in Cali8 with coverage of ~3.5k were identified as viral regions per the criteria outlined in the methods, and confirmed by chromosomal placement in the Nanopore assembly; the contigs with ~2k coverage are mitochondrial. Right panel: Normalized coverage shown as a rolling average across a non-viral contig (upper) and a viral contig (lower) in three distinct Illumina assemblies of *A. arbusculus* Cali8. Cali8-parental was the originally sequenced strain. Cali8-A refers to a strain with abnormal morphology, and Cali8-N refers to a strain presenting normal morphology, as described in the text.

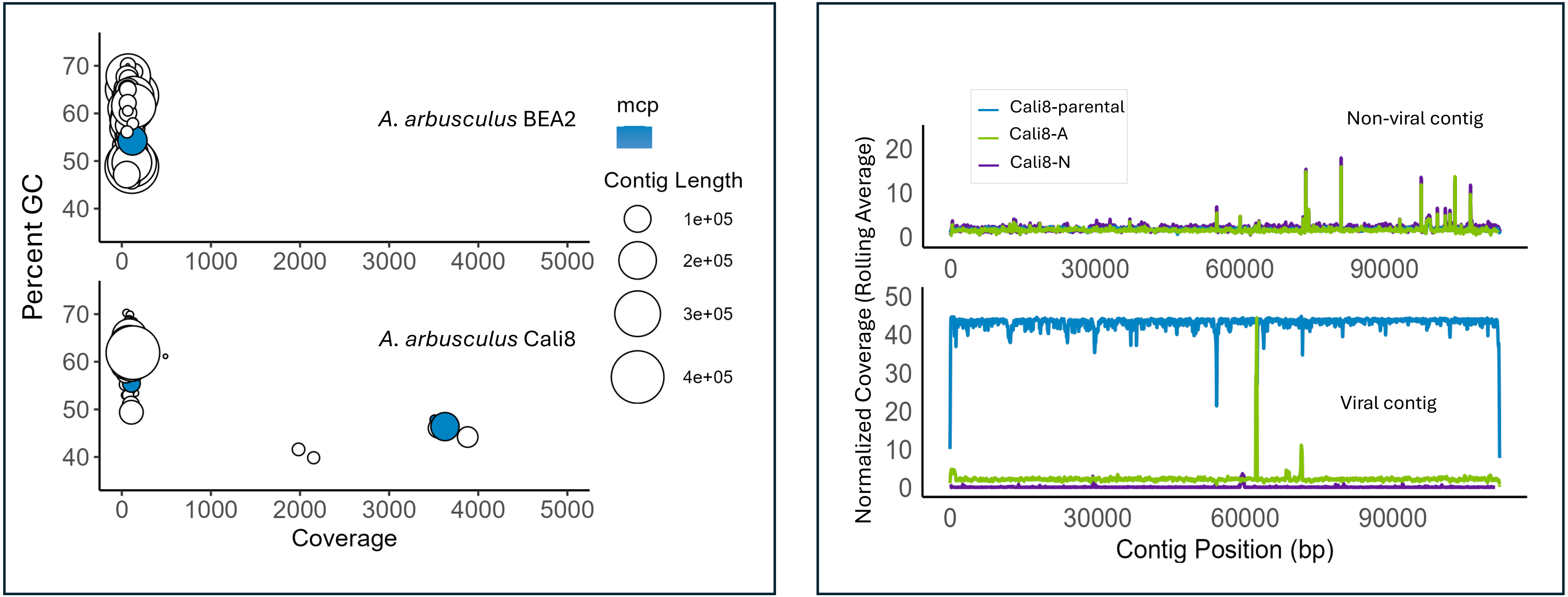

Figure S5. Life cycle of *Allomyces* alternates between diploid sporophyte and haploid gametophyte.

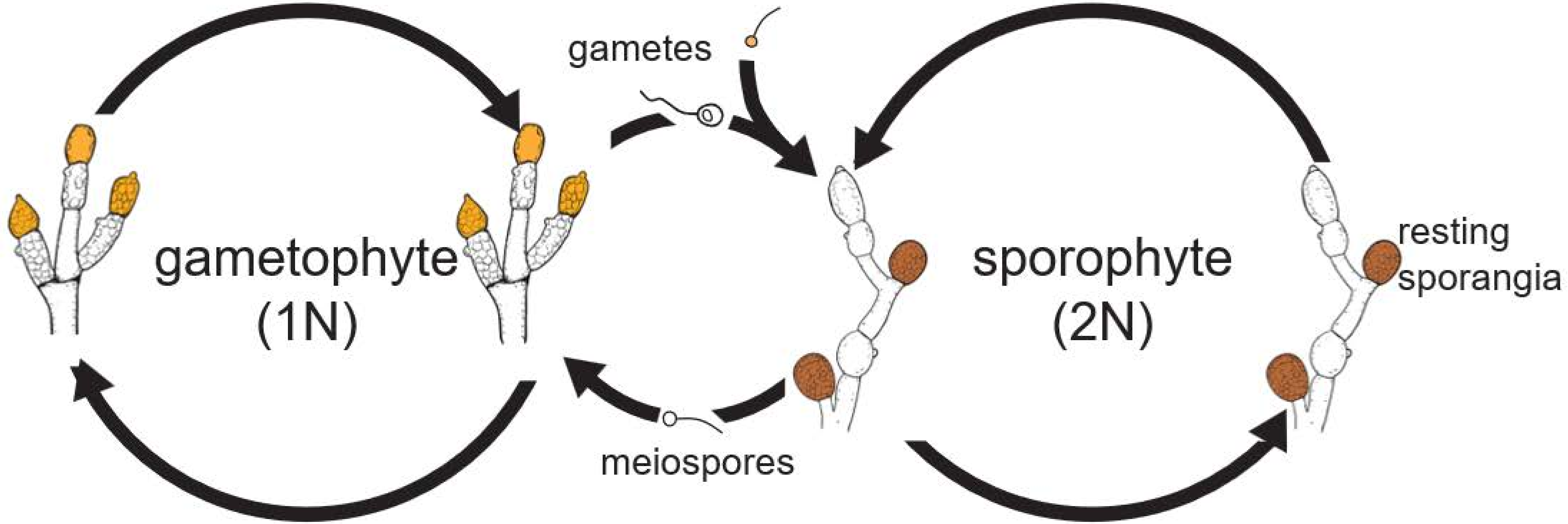

Fig S6: Maximum-likelihood phylogeny of the DNA PolB gene including fungal and *Nucleocytoviricota* reference genes, those from *Mycodnaviridae*, and those previously ascribed to fungal EVEs, with an animal PolB outgroup. Nodes with circles have at least 70% bootstrap support. *Nucleocytoviricota* reference sequences form a well-supported clade that includes the *Mycodnaviridae* and other, distinct, clades of fungal EVEs, suggesting a rich history of association between fungi and giant viruses.

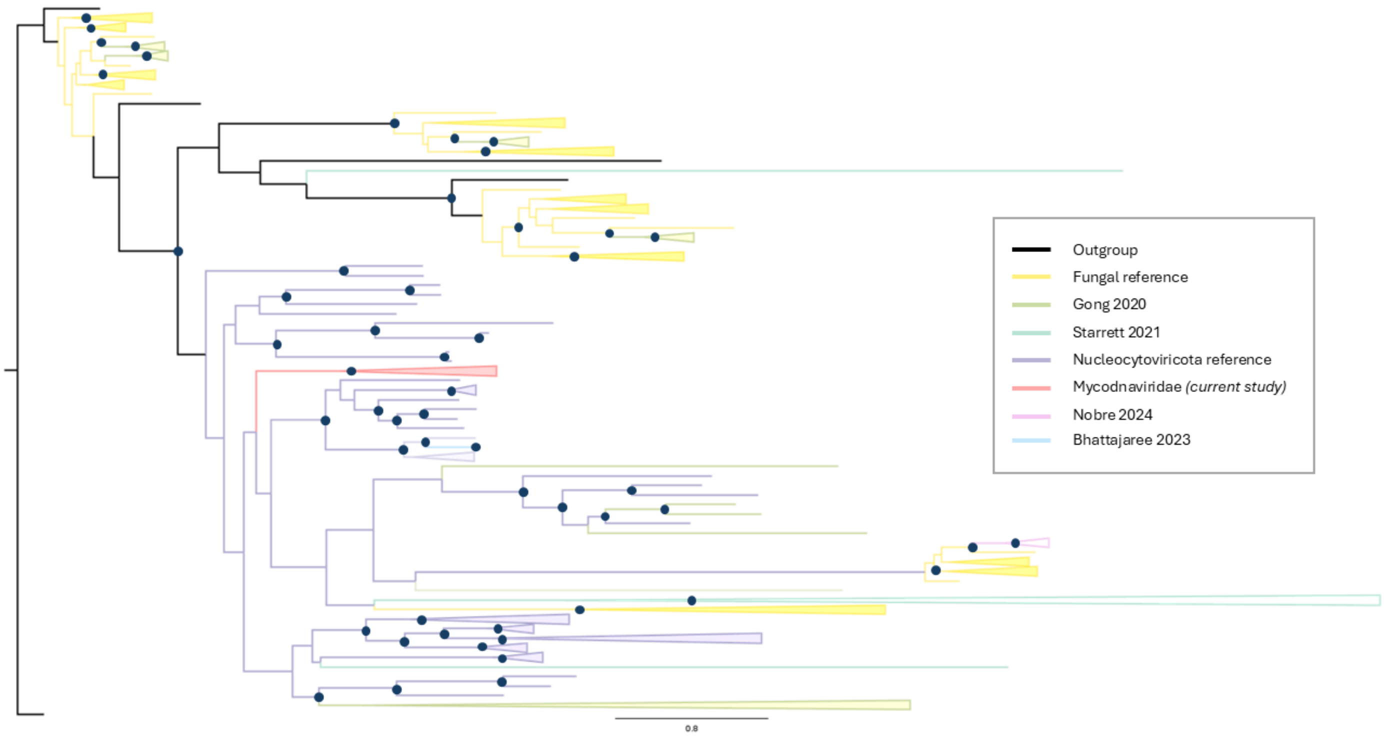

Table S2. Quantitative PCR primer and probe sequences.

| Primer/Probe | Sequence |
|---|---|
| Cali8_cap_44-F | 5'- CCCTCTATGTCCCTCTCATCTT |
| Cali8_cap_44-R | 5'- CGGAACTGGCGGAATGTAAT |
| Cali8_cap_probe | 5'- ATTGCCCTCCTCCACCACGA |
| Cali8_cap_gblock | 5'CAACCCTCTATGTCCCTCTCATCTTCTGGTTCAACCGCAACCCTGGCCTCGCCCTTCCCC TCATTGCCCTCCTCCACCACGACGTCAAAATTAACATTACATTCCGCCAGTTCCGCGACTGC TATGTGCA |
| Cali8_actin_4206-F | 5'- GTAGCAGAGCTTCTCCTTGATG |
| Cali8_actin_4206-R | 5'- GCACTGGCTGCAGAAGAT |
| Cali8_actin_4206_probe | 5'- ACGATTTCGAGCTCGGCCGT |
| Cali8_actin_4206_gblock | 5'TCGAGCGCAACGTAGCAGAGCTTCTCCTTGATGTCGCGCACGATTTCGAGCTCGGCCGT GGTCGTGAGCGAGTGGCCGCGCTCCATGAGGATCTTCTGCAGCCAGTGCGTCAGGTCGCG ACCGGCGAGGTC |
| Cali8_actin PRB Set 1 | 5'-HEX /ACGATTTCG /ZEN /AGCTCGGCCGT /3' IABkFQ/ |
| Cali8_capsid PRB Set 1 | 5'-6-FAM /ATTGCCCTC /ZEN /CTCCACCACGA /3' IABkFQ/ |
